# Unique structural features of the mitochondrial AAA+ protease AFG3L2 reveal the molecular basis for activity in health and disease

**DOI:** 10.1101/551085

**Authors:** Cristina Puchades, Bojian Ding, Albert Song, R. Luke Wiseman, Gabriel C. Lander, Steven E. Glynn

## Abstract

Mitochondrial AAA+ quality control proteases regulate diverse aspects of mitochondrial biology through specialized protein degradation, but the underlying molecular mechanisms that define the diverse activities of these enzymes remain poorly defined. The mitochondrial AAA+ protease AFG3L2 is of particular interest, as genetic mutations localized throughout *AFG3L2* are linked to diverse neurodegenerative disorders. However, a lack of structural data has limited our understanding of how mutations impact enzymatic activity. Here, we used cryo-EM to determine a substrate-bound structure of the catalytic core of human AFG3L2. This structure identifies multiple specialized structural features within AFG3L2 that integrate with conserved structural motifs required for hand-over-hand ATP-dependent substrate translocation to engage, unfold and degrade targeted proteins. Mapping disease-relevant AFG3L2 mutations onto our structure demonstrates that many of these mutations localize to these unique structural features of AFG3L2 and distinctly influence its activity and stability. Our results provide a molecular basis for neurological phenotypes associated with different *AFG3L2* mutations, and establish a structural framework to understand how different members of the AAA+ superfamily achieve specialized, diverse biological functions.

## Introduction

Protein quality control is central to cellular function and survival in all living organisms^1^. Maintaining protein quality control relies on a complex network of ATP-dependent proteases that regulate protein homeostasis (or proteostasis)^2^. The membrane-anchored AAA+ protease FtsH is essential for growth in *E. coli*, and is evolutionarily conserved across all kingdoms of life^3^. These FtsH-related proteins specialize in protein quality control in membranous environments and are found in eukaryotic mitochondrial and chloroplastic membranes, functioning as hexameric, ATP-driven protein quality control systems that process membrane-embedded and associated protein substrates^4,5^. All FtsH-related AAA+ proteases are characterized by a shared topology comprising an amino- (N-)terminal domain, a transmembrane region, an AAA+ ATPase domain, and an M41 metalloprotease domain^6,7^. Notably, mitochondria contain two types of FtsH-related AAA+ proteases, referred to as m- and i-AAA proteases, which are tethered to the mitochondrial inner membrane (IM), but expose their enzymatic domains to the matrix and intermembrane spaces (IMS), respectively^4^ (Figure 1A). We recently solved a cryo-electron microscopy (cryo-EM) structure of the yeast i-AAA YME1, which revealed how a sequential ATP hydrolysis cycle drives a hand-over-hand mechanism of substrate translocation through the central pore of the hexamer^8^. Given the strong structural commonalities shared by all FtsH-related AAA+ proteases, the basic translocation mechanism we described in YME1 is likely to be evolutionarily conserved from bacteria to humans. However, despite this conserved mechanism, FtsH-related proteins have each evolved the ability to recognize and degrade distinct substrates within unique cellular environments, giving rise to this family’s modulation of diverse biological pathways across all kingdoms of life^9–11^. While their distinct substrate specificities likely involve the integration of specialized mechanisms to recognize, recruit, engage, and degrade distinct protein targets in different environments, the molecular basis for this specialization remains poorly understood.

**Figure 1.**
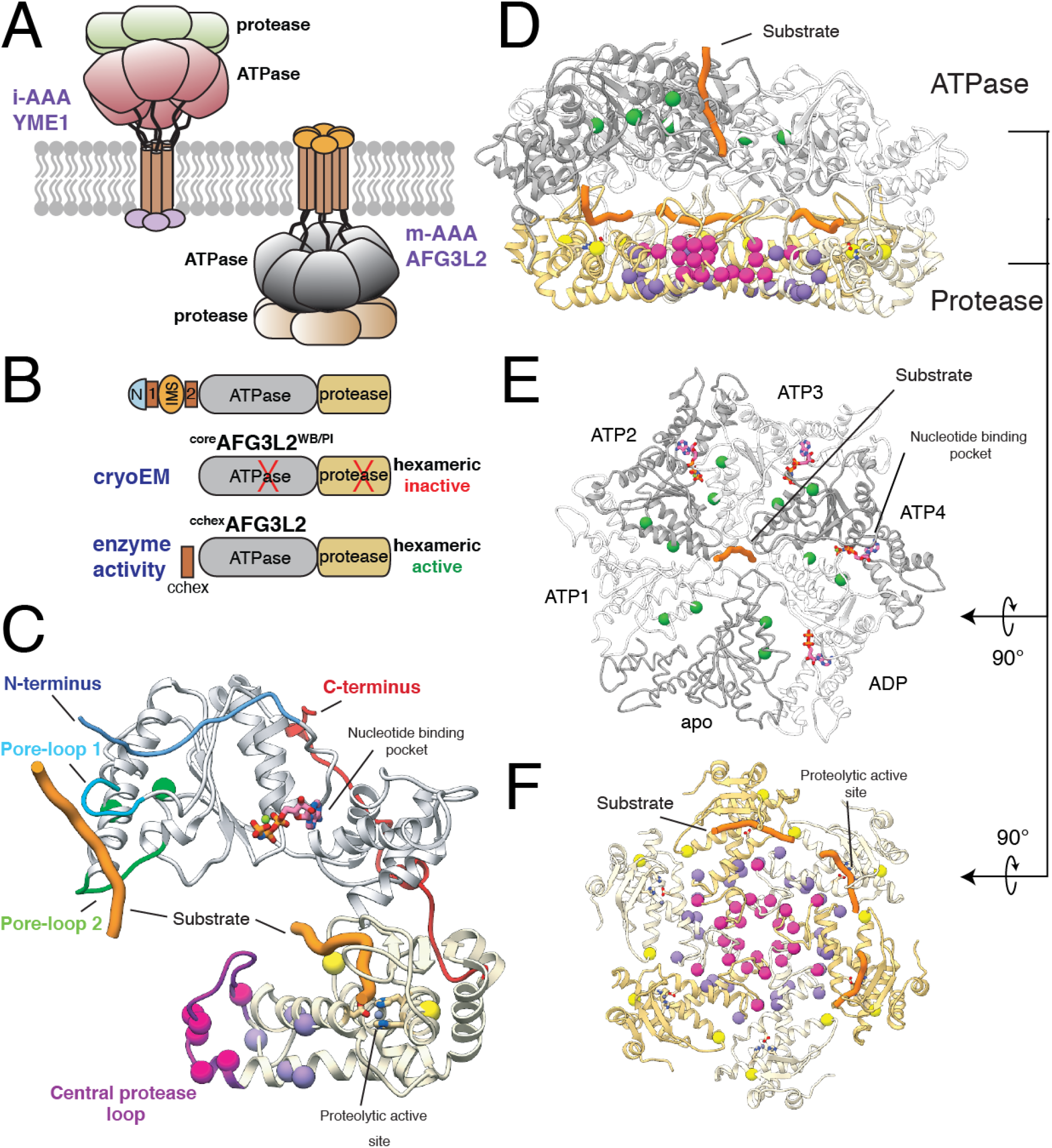
Structure of substrate-bound AFG3L2 reveals four distinct mutational hotspots linked to disease. **A.** Cartoon representation of i- and m-AAA proteases of the mitochondrial inner membrane with the ATPase domains of AFG3L2 colored grey and the M41 zinc metallopeptidase shown in yellow. **B.** Schematic representation of the constructs of the catalytic core of AFG3L2 used in this study. **C.** The atomic model of an AFG3L2 protomer with the ATPase and protease domains colored grey and yellow, respectively. Each residue associated with disease is shown as a sphere colored green, pink, purple, or yellow in accordance with the mutational hotspots identified in our structure. Substrate (orange) interacts with pore-loops 1 (cyan) and 2 (green) of the ATPases, and the proteolytic active site of the protease. The N- and C-termini of the catalytic core are colored blue and red, respectively. **D.** Cutaway side-view of the AFG3L2 homohexamer showing the substrate (orange) in both the ATPase central pore and the proteolytic active sites. The residues linked to disease are shown as spheres colored as in (C). **E.** Top view of the ATPase domains with the disease-related residues depicted as green spheres, demonstrating their proximity to nucleotide (pink) at the intersubunit interface. The two ATPase domains that were rigid body fit into the density are represented as a thin ribbon. **F.** Axial view of the protease ring shows that SCA28 mutations (pink and purple spheres) localize to the intersubunit interfaces and the recessively inherited mutations (yellow spheres) are found at the periphery, in close proximity to the proteolytic active site.

The human m-AAA protease complex assembles as homo-hexamers of AFG3L2 subunits or heterohexamers comprising AFG3L2 subunits and subunits of the closely related homolog paraplegin (SPG7)^12,13^. AFG3L2-containing complexes regulate diverse biological functions, including mitochondrial ribosome assembly, the expression, maturation, and degradation of electron transport chain complexes, supervision of mitochondrial dynamics, and the regulation of calcium homeostasis^14–17^. Furthermore, m-AAA proteases are essential for axonal development in mammals, and loss or reduction of wild-type AFG3L2 lead to pleiotropic phenotypes, such as mitochondrial transport defects, mitochondrial fragmentation, and reductions in both mitochondrial membrane potential and ATP-linked respiration^18–23^. In fact, single point mutations localized throughout the catalytic core of *AFG3L2* are linked to multiple neurodegenerative disorders in humans that present with diverse pathologies and severity. The majority of identified *AFG3L2* mutations are implicated in the autosomal dominant disease spinocerebellar ataxia type 28 (SCA28), primarily characterized by unbalanced standing, progressive gait and limb ataxia, and dysarthria, caused by degeneration of the cerebellum and its afferent and efferent connections^24–32^. An alternative heterozygous *AFG3L2* point mutation has been causatively linked to a disorder called dominant optic atrophy (DOA), whereas homozygous individuals carrying rare mutations in *AFG3L2* present neurodegenerative phenotypes distinct from those associated with either DOA or SCA28^33–36^ (Table S1). The evident genetic relationship between diverse *AFG3L2* mutations and distinct disease severity and neurodegenerative pathologies indicates that these mutations differentially impact m-AAA protease biological activity. Despite this relationship, a lack of structural information has limited our understanding of the molecular mechanisms linking specific mutations to altered enzymatic activity and ultimately pathology.

Here, we present an atomic model of the human AFG3L2 homohexameric m-AAA protease trapped in the act of processing substrate, revealing the network of molecular interactions responsible for substrate engagement, unfolding, transfer across the proteolytic chamber, and proteolysis. While this structure confirms that the fundamental allosteric mechanism relating ATP hydrolysis to substrate translocation is conserved between i-AAA and m-AAAs across eukaryotes, this study also reveals how AFG3L2 has evolved unique structural features for processing substrates located within or in proximity to the matrix-facing surface of the mitochondrial IM. Further, our structure defines how disease-relevant mutations implicated in diverse AFG3L2-associated neurodegenerative disorders distinctly impact the mechanism of action for this AAA+ protease. Thus, our structure-function studies not only provide a much-needed structural framework to understand how different AAA+ proteases build upon a common mechanistic core to uniquely process specific substrates, but also reveal a mechanistic basis to define the distinct neuropathologies associated with different *AFG3L2* mutations.

## Results

### Structure of the human substrate-bound AFG3L2 catalytic core

We solved a ~3.1 Å resolution cryo-EM structure of a truncated construct of AFG3L2 comprising the ATPase and protease domains (residues 272-797; ^core^AFG3L2) (Figures 1 and S1). The construct was stabilized for structural analysis by incorporating ATPase-inactivating Walker B (WB, E408Q) and protease-inactivating (PI, E575Q) mutations. We vitrified the mutant ^core^AFG3L2^WB/PI^ protein in the presence of saturating concentrations of the non-hydrolyzable ATP analog AMP-PNP. The resulting reconstruction was of sufficient quality to build an atomic model of the six protease domains and four of the six ATPase domains (Figures 1 and S1). The large and small ATPase subdomains of one of the well-resolved subunits were rigid-body fit into the low resolution cryo-EM density of the remaining two ATPase domains, providing an atomic model of the entire homohexameric AFG3L2 complex (Figures 1, S2, and Table S2).

Our reconstruction shows that AFG3L2 assembles into two stacked rings with a planar C6 symmetric protease base topped by an asymmetric ATPase spiral (Figures 1D, 1E, and 1F) – a quaternary structure similar to that observed for substrate-bound YME1^8^ (Figure S2A). The three upper subunits of the AFG3L2 staircase are bound to AMP-PNP (termed ATP2-ATP4 for simplicity), while the lowermost subunit is bound to ADP (Figure S2). The limited resolution of the remaining two subunits did not allow for unambiguous identification of nucleotide state, although the positioning of these subunits within the staircase correspond to the nucleotide-free (apo) ‘step’-subunit and the uppermost ATP-bound subunit (ATP1) within YME1 (Figure S2A). Importantly, we observe density corresponding to substrate peptide threaded through the center of the ATPase spiral of AFG3L2, as has been observed for other AAA+ proteins including YME1^8,37–41^. Unexpectedly and in contrast to YME1, this translocating substrate density directly contacts the center of the protease ring. Furthermore, use of the protease inactivating (E575Q, denoted AFG3L2^PI^) mutation resulted in observable density corresponding to substrate trapped in the proteolytic active sites of the ATP2-4 subunits (Figure 1C and 1F). Thus, our AFG3L2 structure provides a unique opportunity to dissect the complete mechanism of substrate translocation and degradation by this AAA+ protease, and its relation to disease.

### Modulation of a conserved translocation mechanism by AFG3L2-specific pore-loop 1 residues

Numerous structures of substrate-bound AAA+ unfoldases have been recently solved by cryo-EM, and suggest conservation of a mechanism for substrate translocation wherein a sequential ATP hydrolysis cycle powers a hand-over-hand conveyance of an unfolded polypeptide through the central channel of the hexamer^8,37–41^. Like in other AAA+ ATPases, an aromatic residue within the ATPase domain pore-loop 1 of AFG3L2 (F381) forms a spiral staircase around the translocating substrate, and intercalates into its backbone (Figures 2A, 2C, S2B, and S2C). To verify the crucial role of F381 in substrate translocation, we used established assays to monitor ATP hydrolysis and substrate degradation for an engineered hexameric AFG3L2, where the transmembrane domains were substituted by a hexamerizing coiled coil (^cchex^AFG3L2)^42^ (Figure 1B). As previously shown, the coiled coil ensures hexamerization of active subunits in the absence of any stabilizing mutations^43,44^. Incorporation of an F381A substitution abolished substrate degradation, while only mildly impacting ATP hydrolysis (Figure 2B), confirming that the aromatic pore-loop 1 residue in AFG3L2 plays a conserved, essential role in substrate translocation.

**Figure 2.**
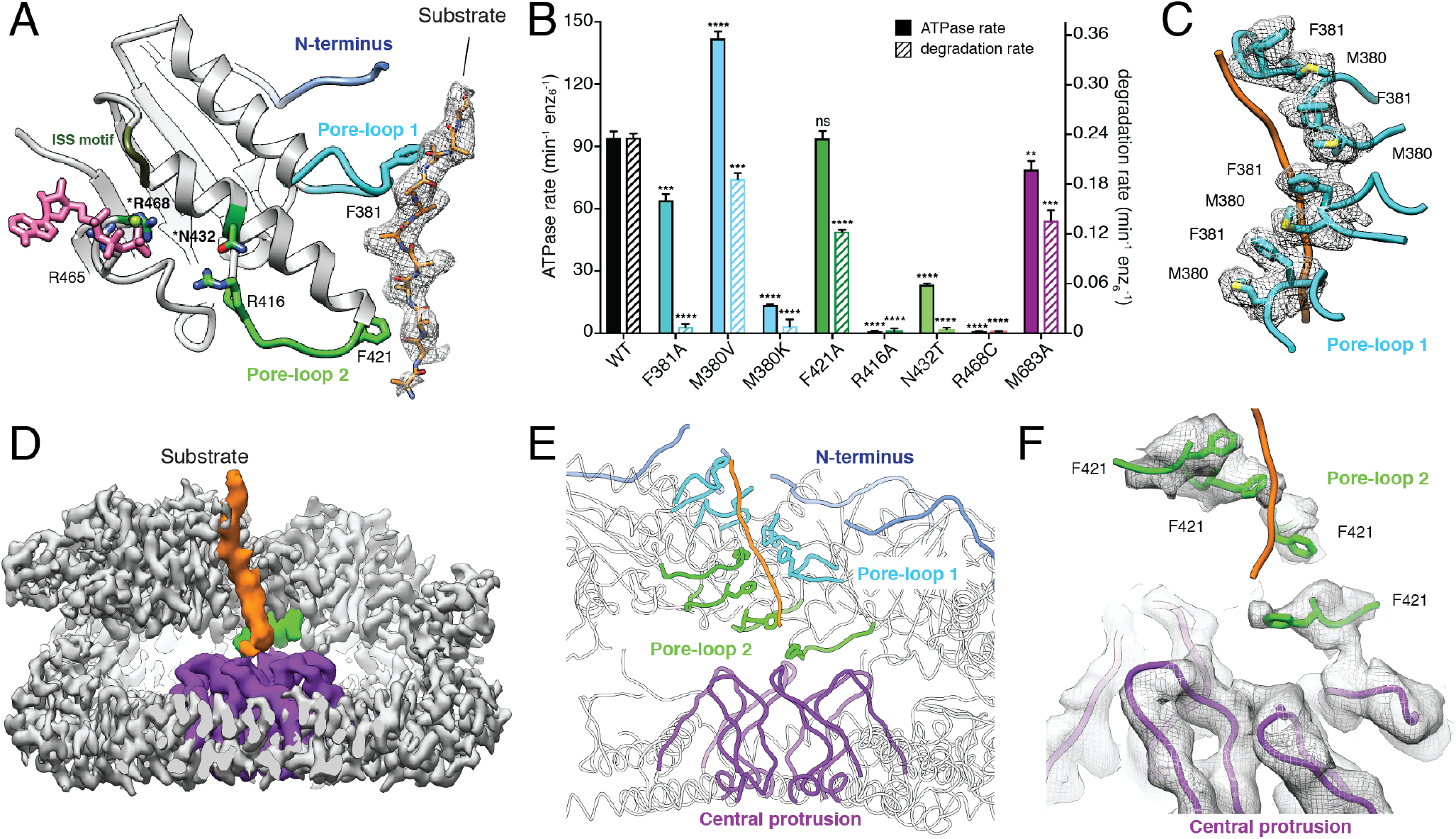
Unique features of AFG3L2 mediate substrate transfer from the ATPase to the protease. **A.** The ATPase domain’s N-terminus (top, blue), pore-loop 1 (middle, cyan), and pore-loop 2 (bottom, green) directly contact the trapped substrate polypeptide (cryo-EM density shown as mesh). The nucleotide from the adjacent subunit is shown (pink), highlighting the position of the disease-associated residues R468 and N432 relative to the neighboring subunit’s nucleotide-binding pocket. **B.** ATPase and substrate degradation rates for ^cchex^AFG3L2 and its variants with single-point substitutions in pore-loops 1, 2, and the central protrusion validate the functional relevance of each of these regions for substrate processing. Values are means of independent replicates (n ≥ 3) ± SD. **P ≤ 0.01, ***P ≤ 0.001, ****P ≤ 0.0001 as calculated using Student’s two-tailed t test and statistical significances are shown relative to WT ^cchex^AFG3L2. **C.** Details of the pore-loop 1 spiral staircase with residues F381 and M380 illustrated as sticks and the cryo-EM density shown as a grey mesh. M380 interacts with F381 of the adjacent subunit, forming a continuous network that surrounds the substrate (orange). **D.** Cutaway view of the AFG3L2 homohexamer cryo-EM reconstruction shows that the pore-loop 2 of the lowest subunit (green) contacts the central protease protrusion (purple). The unsharpened density for the substrate is shown in orange, demonstrating direct contacts with both pore-loop 2 and the central protrusion. **E.** Ribbon representation of the substrate peptide (orange) in the central pore, with pore-loops 1 and 2 colored blue and green, respectively, to highlight the two staircases that simultaneously interact with the substrate. The lowest pore-loop 2 directly contacts the central protrusion of the protease domains, shown in purple. **F.** Detailed view of the staircase formed by residues F421 from pore-loop 2 (green) and contact with the central protrusion (purple). Modeled substrate is orange, and cryo-EM density is shown as a mesh.

A key requirement for the ATP-driven hand-over-hand substrate translocation mechanism is allosteric transmission of nucleotide state to the pore-loops. This allostery was previously described in YME1, wherein consecutive Asp-Gly-Phe residues located at the interface between neighboring ATPases alternates conformations between a compact α-helix and an extended loop that spans the nucleotide-binding pocket at the inter-protomer interface^8^. Conformational switching of this region, termed the inter-subunit signaling (ISS) motif, is directly controlled by the nucleotide state, which in turn allosterically defines the positions of pore-loops 1 and 2^8,45^. Together, this gives rise to a mechanism wherein the pore loops of ATP-bound subunits engage the substrate, ATP hydrolysis in the lowermost subunit causes its pore loops to detach from the substrate, and nucleotide exchange enables re-engagement of substrate at the top of the staircase. The correlation between nucleotide state, conformation of the ISS and positioning of the pore loops is conserved in our AFG3L2 structure, indicating that allosteric regulation of pore-loop conformation in response to nucleotide state, and thereby the fundamental mechanism of substrate translocation, is conserved between i- and m-AAA proteases across eukaryotes (Figure S2B and S2C).

Despite conservation of the overall mechanism, the pore-loops of AFG3L2 contain structural features distinct from those observed in YME1 or other AAA+ ATPases, which integrate with the core components to modulate substrate translocation. For example, the methionine residue immediately preceding the F381 pore-loop 1 aromatic residue, M380, contacts F381 of the counterclockwise neighboring subunit to form a continuous chain of residues that wrap around the substrate in the central channel (Figure 2C), an organization distinct from that observed in YME1. This methionine residue is largely conserved in FtsH-related enzymes, with the notable exception of YME1, but is highly variable across the broader AAA+ protease family (Figure S3A). An M380V mutation, which converts this methionine to the corresponding valine residue seen in YME1, increased the ATP hydrolysis rate while moderately decreasing substrate degradation, indicating reduced enzymatic efficiency (Figure 2B). Moreover, replacing this methionine with a lysine residue observed in five of the six 26S proteasome ATPase subunits (M380K) completely abolished AFG3L2 activity, consistent with previous reports showing that the corresponding Met to Lys mutations in the AFG3L2 homologs Yta10 and Yta12 obliterated substrate processing in yeast^46^. Together, our data show that the methionine residue immediately prior to the conserved pore-loop 1 aromatic gives rise to a distinct molecular configuration of the central channel that specifically influences AFG3L2 substrate translocation. Moreover, this aromatic-prior residue likely contributes to the diverse substrate specificities observed across the AAA+ ATPase family by providing a means of fine-tuning the electrostatic and steric characteristics of the substrate-interacting pore-loops.

### Pore-loop 2 coordinates substrate transfer to a central protrusion within the protease domain

Pore-loop 2 is the least conserved region of the ATPase domain across FtsH-related AAA+ proteases. Surprisingly, we observe that a four-residue insertion in pore-loop 2 of AFG3L2 positions the central aromatic residue F421 closer to the substrate than what is observed for YME1^8^, creating a second spiral staircase that directly engages the substrate – an interaction not observed in any other substrate-bound AAA+ ATPase solved to date. Incorporation of an F421A substitution in ^cchex^AFG3L2 impaired substrate degradation with no impact on ATPase activity, confirming an important role for the pore-loop 2 aromatic residue in substrate translocation (Figure 2B). These additional contacts with the translocating substrate likely increase the ATPase’s “grip” on substrate for optimal processing of AFG3L2-specific substrates.

Importantly, the extended AFG3L2 pore-loop 2 positions the F421 residue of the two lowest subunits within the spiral staircase (ATP4 and ADP; Figure 2E and 2F) deep in the interior of the proteolytic cavity. In this configuration, F421 of the ADP subunit’s pore-loop 2 contacts a ‘central protrusion’ within the protease domain. This protrusion is formed by residues 673-695 of each subunit, which form a hexamer of six upward-projecting loops at the central base of the protease ring, directed toward the incoming translocating substrate (Figure 2D and 2E). While sequences corresponding to the central protease loop are present in other AAA+ proteases, such as FtsH (Figure S3B), our structure captures the ordered interaction between the central protrusion and the pore loops for the first time, likely a result of the presence of substrate^47–49^. Six M683 residues crown the top of the central protrusion, and the lowermost F421 residue from pore-loop 2 directly contacts the nearest M683 in a manner that is reminiscent of the Phe-Met interactions observed in the pore-loop 1 spiral (Figure 2D-2F). Moreover, at low sigma values the density corresponding to translocating substrate extends beyond the lowermost F421, towards the center of the protease and in close proximity to the M683 crown (Figure 2D). We incorporated an M683A substitution into ^cchex^AFG3L2 to define the importance of the F421-M683 interaction in substrate processing, and observed moderate reduction in substrate degradation while minimally affecting ATP hydrolysis, mirroring the impact of the F421A mutation (Figure 2B). We thus conclude that the unexpected organization of the extended pore-loop 2 and the central protease loops in AFG3L2 likely plays an important role in facilitating substrate transfer from the ATPase spiral to the proteolytic ring. Notably, both the prolonged pore-loop 2 as well as the extended central protease loop are present in m-AAA proteases but absent in i-AAA proteases across evolution (Figure S3), suggesting that the distinct substrate handling identified here has important implications for the differential functional roles of m- and i-AAA+ proteases.

### The N-terminus of the AFG3L2 ATPase domain mediates membrane-proximal interactions with the substrate

In addition to the distinct pore-loop interactions revealed by our structure, we also observe a previously undescribed organization of the ATPase domain’s N-terminus. This N-terminus connects the catalytic core of AFG3L2 to the transmembrane domains and begins with a flexible glycine-rich region (residues 272-287). Despite this region being present in our ^core^AFG3L2 construct, it was not observed in our cryo-EM reconstruction (Figure S3A). Surprisingly, however, our structure shows that the residues immediately following this region (288-295) form an additional spiral staircase above the ATPase domains that surrounds and contacts the translocating substrate in the central channel (blue in Figure 3A). Interestingly, ^cchex^AFG3L2 constructs containing deletion of the ordered, substrate-interacting ATPase N-terminus (residues 272-295) exhibited significantly impaired substrate degradation without impacting hydrolysis, whereas deletion of just the flexible Gly-rich linker region (residues 272-281) did not influence either AFG3L2 activity. This indicates that interactions between the substrate and the spiral staircase formed by the N-terminal residues of ATPase domain are important for substrate engagement and translocation into the catalytic core of AFG3L2.

After leaving the site of substrate interaction near the central channel, each N-terminus snakes across the interface between neighboring ATPases, directly above the intersubunit interface where the ISS motif engages the clockwise adjacent subunit (Figure 3B and 3C). Interestingly, L299 of each N-terminus inserts into a hydrophobic groove formed by the trans-subunit packing of the central ISS phenylalanine residue against phenylalanine residues within each ATP-bound subunit (Figure 3C). In the ADP-bound subunit, for which there are no ISS interactions and thus the hydrophobic groove is absent, the N-terminus L299 is repositioned closer to the phenylalanine of the cis subunit. The maintenance of these hydrophobic interactions regardless of nucleotide state suggested that the packing of L299 against the ATPase body may stabilize the N-terminus and promote interaction with substrate. As we predicted, ^cchex^AFG3L2 containing the L299A mutation shows 50% reductions in substrate degradation without impacting ATP hydrolysis (Figure 3D). Taken together, this reveals a previously unanticipated role for the AFG3L2 ATPase N-terminus in mediating contacts with the substrate in the membrane-proximal region of the complex.

**Figure 3.**
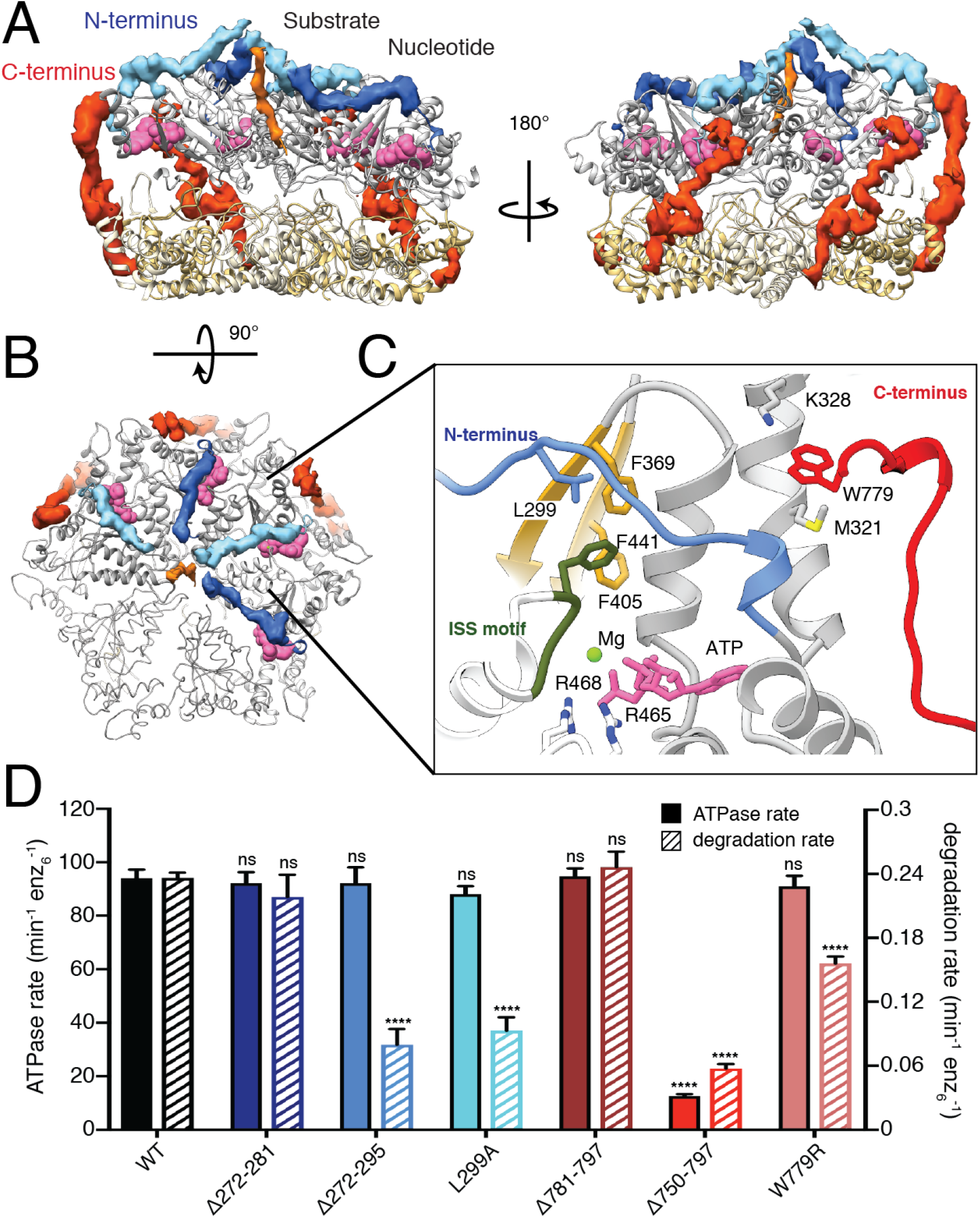
The N- and C-termini of the catalytic core of AFG3L2 play an essential structural role at the membrane proximal region. **A.** Atomic model of the catalytic core of AFG3L2 shown as a ribbon with the cryo-EM density of the N- and C-termini shown in blue and red, respectively. The N-termini of the ATPase domains extend across the nucleotide binding pocket (nucleotides as pink spheres), and form a spiral staircase around the substrate (orange). The C-termini extend across the outer surface of the complex. **B.** Top view of the ATPase staircase shows how the N-termini cross over the nucleotide-binding pocket at the inter-subunit interface and contact the substrate. **C.** Close-up of the interaction network in the AMPPNP-bound nucleotide-binding pocket: 1) The arginine fingers (R465 and R468) of the neighboring subunit (in white) contact the nucleotide, 2) F369 interacts with F441 of the ISS motif (dark green) of the neighboring subunit, 3) The N-terminal L299 residue is positioned above these two F residues, and 4) M321 and K328 in the N-terminus of the ATPase domain interact with W779 in the C-terminus of the protease domain. The two ATPase domains that were rigid body fit into the density are represented as a thin ribbon. **D.** ATPase and substrate degradation rates for ^cchex^AFG3L2 and its variants with truncations or single-point substitutions in the N- and C-termini reveal that the ordered residues in these regions are important for AFG3L2 activity. Single-point mutations in L299 and W779 in the N- and C-termini, respectively, highlight the functional relevance of the interactions mediated by these residues.

### The AFG3L2 C-terminus encircles the hexamer to adopt a membrane proximal position

Our construct contained the complete carboxy- (C-) terminus of AFG3L2, comprising residues 750-797 – a region that is highly variable among FtsH-related AAA+ proteases and, in AFG3L2, contains a number of charged residues at the far C-terminus (Figure S3B). While we do not observe density for the highly charged C-terminal tail in the structure (residues 780-797), well-defined density is present for the residues immediately preceding the tail (residues 750-779) in the three AMP-PNP bound subunits (ATP2-4) (Figure 3A). Unexpectedly, this ordered C-terminal sequence extends upwards from the base of the protease domain along the exterior surface of the complex to interact with both the protease and the ATPase domains of the cis subunit, and the interdomain linker of the counterclockwise adjacent subunit. Interestingly, truncation of the C-terminus at residue 750 in the ^core^AFG3L2^WB/PI^ construct, which is not stabilized by the hexamerizing coiled coil domain, dramatically decreased recovery of assembled AFG3L2 hexamers as measured by size exclusion chromatography (SEC; Figure S4A). In contrast, removal of the unobserved charged tail alone (residues 781-797) did not impact complex recovery. This indicates that the interaction of the C-terminal residues 750-779 across neighboring ATPase domains is important for stabilization and assembly of the active hexamer. Consistent with this, deletion of residues 750-797 of ^cchex^AFG3L2, where hexamers are held together by coiled-coil domains, reduced both ATPase activity and substrate degradation, whereas deletion of residues 781-797 had no significant effect on either activity. Although the highly charged C-terminal tail is not visible in our structure, the unexpected arrangement of the remainder of the C-terminus positions the tail at the membrane-proximal face of AFG3L2. The final C-terminal residue for which we observe well-ordered density in our structure is residue W779, which is sandwiched between M321 and K328 at the membrane-proximal surface of the ATPase domain (Figure 3C). These stabilizing sulfur-aromatic and cation-π interactions likely secure this tryptophan residue in position, anchoring the base of the charged C-terminal tail at the membrane-proximal surface of AFG3L2 for potential substrate interaction. In support of this, incorporating a W779R substitution into^cchex^AFG3L2 impaired substrate degradation without impacting ATP hydrolysis (Figure 3D). Thus, our results reveal an important role for the C-terminus in dictating complex stability, and show that interactions driven by the organization of C-terminal residues influence important aspects of AFG3L2 substrate processing.

### Disease-relevant AFG3L2 mutations localize to four ‘hotspots’ on the AFG3L2 structure

Our structure reveals how structural features unique to AFG3L2 integrate with a core nucleotide-driven allosteric mechanism to drive substrate translocation. Thus, we next sought to evaluate how specific disease-relevant AFG3L2 mutations impact its activity. Mapping disease-relevant AFG3L2 mutations onto our structure reveals that all mutations currently linked to disease localize to four distinct ‘hotspots’ within the hexameric complex (Figure 1C-1F). Intriguingly, all of the autosomal dominant mutations localize to intersubunit interfaces of the hexamer, including 1) the lateral interface between adjacent ATPase subunits where the nucleotide binding pocket is formed, 2) the tightly interconnected helices that dominate the lateral interface of the protease domain, and 3) the central protrusion of the AFG3L2 protease ring that is involved in substrate transfer to the protease domains. In contrast, recessive mutations localize near the protease active sites in the periphery of the protease ring. The clustering of these mutations into distinct regions in the structure suggests distinct effects on AFG3L2 activity.

### Disease mutations in the AAA+ domains disrupt nucleotide-dependent substrate translocation

Two disease-relevant mutations, N432T (linked to SCA28) and R468C (associated with DOA), localize to the vicinity of the nucleotide-binding pocket at the ATPase intersubunit interface^25,33,34^. Our structure shows that R468 is one of the two arginine finger residues of AFG3L2 that projects from the adjacent large subdomain to coordinate interactions with the γ-phosphate in the adjacent subunit (Figure 3C and S2B). These arginine finger residues are highly conserved across classical AAA+ enzymes and are suggested to play roles in ATP binding and/or intersubunit interactions^50^. In fact, mutation of analogous arginine fingers in bacterial FtsH obliterated activity^51^. Accordingly, the R468C patient mutation completely abolishes ATP hydrolysis and subsequent substrate degradation when incorporated into ^cchex^AFG3L2 (Figure 4C). Furthermore, and consistent with impaired ATP binding at the intersubunit interface, R468C decreased recovery of AFG3L2 hexamers by SEC, closely mirroring the behavior of the ATP binding incompetent control, Walker A (K354A, denoted AFG3L2^wa^) (Figure S4B).

**Figure 4.**
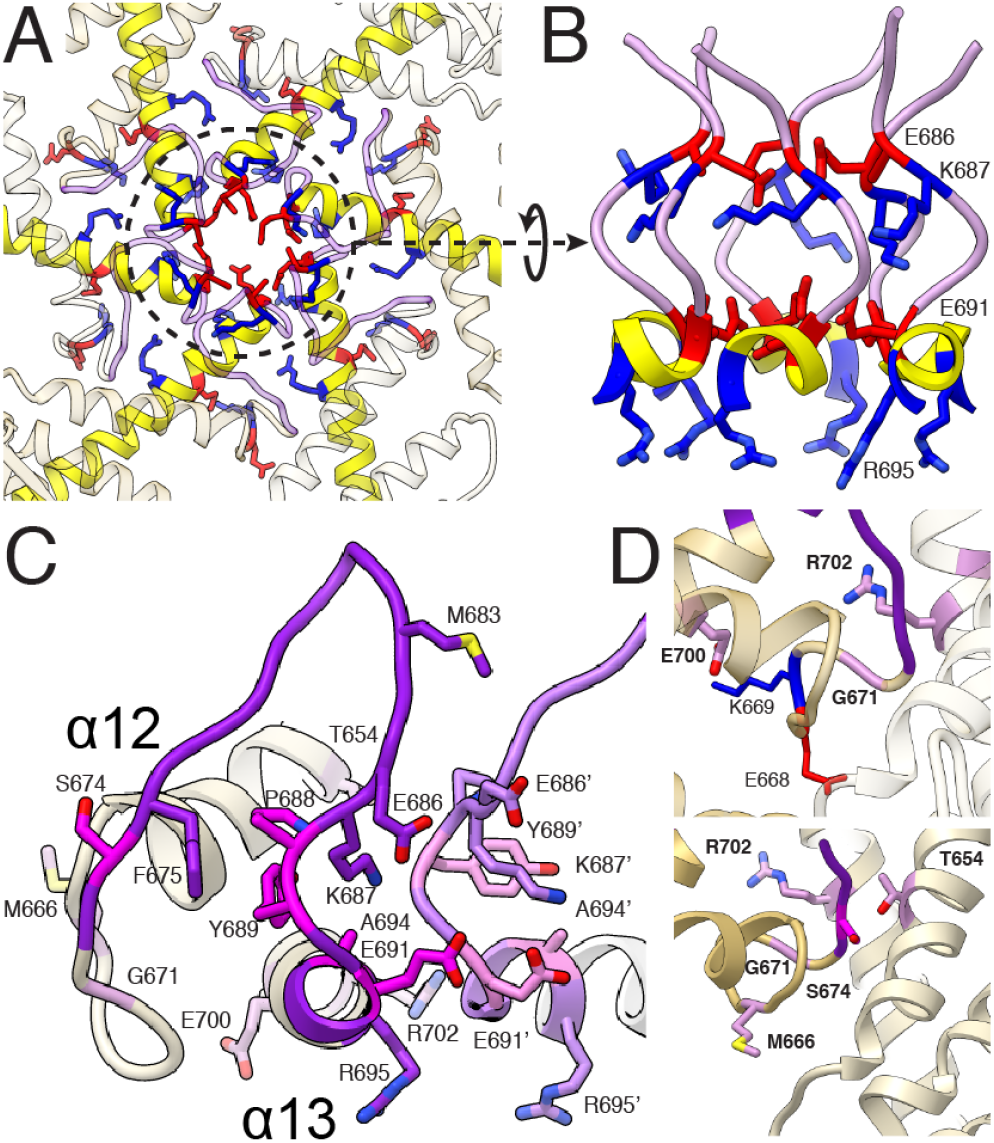
SCA28-associated residues of the protease domain mediate important inter-subunit contacts. **A.** Axial view of the center of the protease ring with the central protrusion colored light purple, and the negatively and positively charged residues depicted as red and blue sticks, respectively. **B.** Side view of the central protrusion, colored as in (A), reveals stacked rings of alternating charge that face the central channel. **C.** A close-up of residues 653-703, where all residues linked to SCA28 in the protease domains are found, shows the tightly packed inter- and intra-subunit interaction network at the center of the protease ring. Disease-associated residues are shown as sticks and highlighted in magenta. **D.** Close up views of SCA28-associated residues at the base of the central protrusion. Top, close-up of the charge cluster at the lateral interface of the protease domains. Disease-related G671 (light purple) is found at the base of the central protrusion (dark purple) in a loop that inserts between helices α13 of neighboring subunits. On the same loop, E668 (red) and K669 (blue) interact with disease-associated residues E700 and R702 on helix α13 (light purple). Bottom, close-up of the interactions between adjacent α12 helices and the central protrusion. S674 (magenta) from the central protease loop packs against T654 (light purple) on helix α12 of the neighboring subunit, and mutations in either of these residues are linked to disease. SCA28-associated residue M666 (light purple) is found at the interface between adjacent α12 helices.

Previously, homology models of FtsH were used to predict that the N432T mutation might directly impair substrate engagement in the central pore^25^. Our structure shows that while N432 is not part of the central pore of the ATPases, N432 interacts with R416 within the extended pore-loop 2, thereby positioning this loop to facilitate substrate engagement (Figure 2A). Both N432 and R416 are strictly conserved within FtsH family, implying that the interaction we observe in AFG3L2 exists in other family members (Figure S3A). In agreement with a key functional role for the N432-R416 interaction, both the R416A and disease-relevant N432T substitutions completely abolished substrate degradation in^cchex^ AFG3L2. Surprisingly, we found that these two mutations also significantly reduced ATPase activity. The decreased ATP hydrolysis rate for N432T can be explained by its localization within helix α5 immediately adjacent to the ISS motif, suggesting that mutation of this residue might disrupt allosteric coordination of ATP hydrolysis. In agreement, this mutant displays a significantly reduced maximal ATPase rate but retains high affinity for ATP in Michaelis-Menten kinetic measurements, indicating that N432T impairs catalysis but does not interfere with nucleotide binding (Figure S4F). Thus, our structure-function studies indicate that the disease-relevant N432T substitution alters AFG3L2 activity through multiple mechanisms, allosterically affecting both, the ATP hydrolysis cycle and pore-loop 2 organization.

### Disease mutations in the protease domain concentrate near the central protrusion

With the exception of N432T, all SCA28-associated mutations identified to date localize to two specific hotspots within or at the base of the central protease loop in the AFG3L2 protease domain (Figure 4A-4D). As discussed above, the central protease loops form a hexameric channel-like protrusion at the center of the proteolytic ring, which interacts with pore-loop 2 of the ATPases and the translocating substrate (Figure 2D-2F). In each subunit, the base of the central protease loop is stabilized by π-stacking intrasubunit interactions between Y689 and F675 (Figure 4C). Intriguingly, layered rings alternating between acidic (E686 and E691) and basic (K687 and R695) residues create a highly charged channel surface (Figure 4A and 4B). All of the mutations localized to this region line the channel of the central protrusion and likely impact the integrity of its structure. E691K swaps the charge of one of the acidic rings, which would disrupt the integrated charged ring system we observe in the channel (Figure 4B). Similarly, A694E results in a negatively charged residue being positioned adjacent to E691, introducing a charge repulsion between these residues (Figure 4B). At the base of the central protease loop, substitutions replacing Y689 (Y689N and Y689H) would eliminate the π-stacking interactions that maintain loop integrity (Figure 4C). Finally, P688 introduces a sharp turn at the base of the loop that correctly positions Y689 and K687 for interaction with other residues in this region (Figure 4C). The P688T substitution will likely add flexibility at this position that will decrease the stability of the central protrusion.

In contrast to the central protease loop, residues at the base of the central protrusion are highly conserved across FtsH-related proteases, and our structure indicates that the majority of these residues are involved in intersubunit interactions that stabilize the proteolytic ring. Notably, all remaining SCA28-associated residues are located in this region, including one of the most highly mutated residues in disease, M666, which integrates into a tightly packed hydrophobic pocket at the intersubunit interface (Figure 4D). Mutation of M666 to Thr or Val would perturb this hydrophobic pocket. Importantly, mutation of M666 to Arg, which would severely disrupt this pocket by introducing a positively charged residue, results in one of the most severe disease phenotypes in patients. Substitutions R702Q and G671R/E would destabilize a charged cluster at the base of the central protrusion (Figure 4D). Similarly, the E700K mutation disrupts anion-cation interactions with K669 at the base of the central protrusion in the same subunit (Figure 4D). Disease states brought about by destabilization of the protease domain can also be ascribed to mutation of the polar residues T654 and S674 to bulky, hydrophobic residues (T654I and S674L), which would introduce steric clashes into otherwise closely packed intersubunit interactions (Figure 4D). Based on these structural observations, we predict that all SCA28-associated mutations localized within or at the base of the central protrusion destabilize the protease, thereby impacting AFG3L2 function.

Consistent with this, incorporating these protease mutations into ^core^AFG3L32^WB/PI^ significantly impaired recovery of the hexameric protein by SEC, albeit to different extents (Figure S4C and S4D). For example, the M666R mutant, which disrupts the hydrophobic interface between protease subunits, completely abolished hexamer recovery, while the P688T mutant that increases flexibility at the base of the protrusion only modestly affected stability (Figure S4C and S4D). This global destabilization of the catalytic core was further evident when we incorporated protease mutations into ^cchex^AFG3L2, where hexamers are held together by coiled coil interactions. Despite hexamer formation, all protease mutations show reductions in both ATP hydrolysis and substrate degradation that reflect the degree of destabilization observed by SEC (Figure 5C). Moreover, reductions in activity *in vitro* appear to correlate with disease severity in patients. For instance, P688T presents only a modest reduction in activity *in vitro*, and is associated with the mildest condition in SCA28 patients, whereas M666R shows a near complete loss of activity and is associated with a very severe, early onset phenotype in SCA28 patients^26,29^.

To further define the impact of these mutations on AFG3L2 stability, we incorporated the mild mutation P688T and the severe mutations M666R and E691K into C-terminally Flag-tagged AFG3L2 (AFG3L2^FT^) and monitored assembly and stability by Blue-Native (BN-PAGE) and SDS-PAGE. Interestingly, all of these protease mutations destabilized AFG3L2 oligomers, as measured by BN-PAGE (Figure 5A). Furthermore, using antibodies that recognize the N- or C-terminus of AFG3L2^FT^, we observed the accumulation of an N-terminal cleavage product of ~50 kDa in SDS-PAGE that corresponds to loss of the C-terminal protease domain (Figure 5B). This accumulation was most evident for M666R and E691K, reflecting the enhanced destabilization of these mutants. These results further substantiate a model whereby mutations in the protease domain destabilize the protease ring, leading to SCA28, with the extent of destabilization being an important factor for disease severity and age of onset in patients (Table S1). Importantly, these results were in stark contrast to those observed when the ATPase mutants N432T and R468C were analyzed using the same approach (Figure 5B and 5C). To determine whether their distinct profiles are related to direct impairment of the ATP hydrolysis cycle, we expressed control mutants Walker A (prevents ATP binding) and Walker B (prevents ATP hydrolysis). As in our SEC experiments, the Walker B mutant increased stability of the AFG3L2 hexamer, whereas in the Walker A mutant we detected the appearance of AFG3L2 monomers, indicating disassembly of the complex. These results indicate that ATP binding is important for the stability of the AFG3L2 hexamer, likely because the nucleotide-binding pocket is formed at the intersubunit interface, and ATP mediates substantial intersubunit contacts. In agreement with our *in vitro* data, R468C closely mirrors the behavior of the ATP binding-incompetent Walker A control, potentially reflecting the unique DOA pathology observed in patients harboring this mutation. While the SDS-PAGE profile of N432T is similar to that of the Walker B mutant, N432T causes a mild destabilization of AFG3L2 oligomers by BNPAGE (Figure 5A). Our biochemical and structural data indicate that the N432T substitution might allosterically impact both ATP hydrolysis and pore-loop conformation (Figure 2). Thus, the decreased stability observed in cells might be due to concomitant impairment of substrate binding, providing a potential explanation for the strong dominant negative effect seen for this mutation^25^. To summarize the above sections, we identify three distinct molecular mechanisms by which autosomal dominant AFG3L2 mutations influence the activity of this AAA+ protease, and these different mechanisms appear to correlate to distinct phenotypes and disease severity in patients (Table S1).

**Figure 5.**
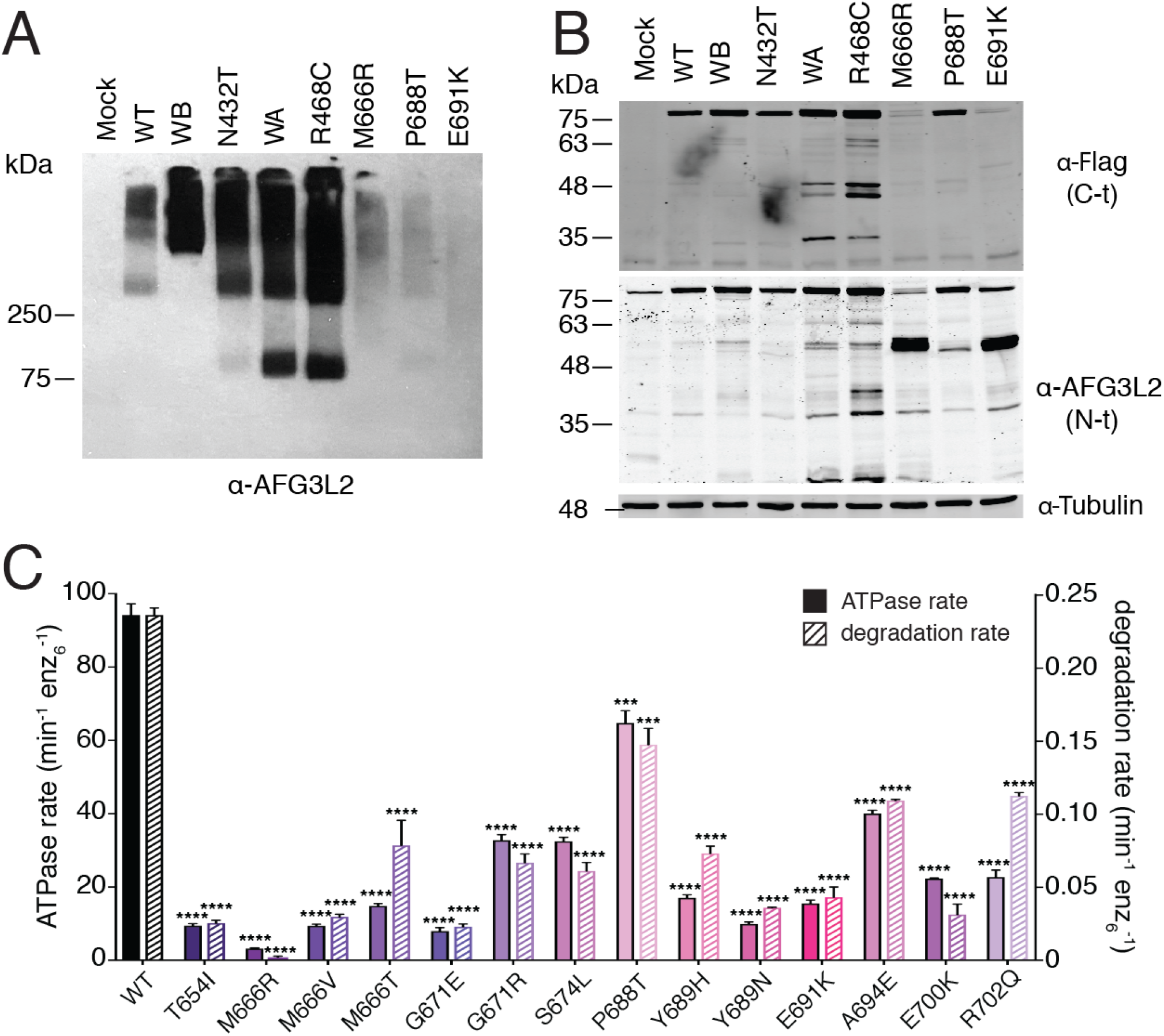
Autosomal dominant mutations impair AFG3L2 function to different extents through three distinct molecular mechanisms. **A.** BNPAGE and **B.** SDS-PAGE analysis of C-terminally Flag-tagged AFG3L2 containing representative disease mutants (N432T, R468C, M666R, P688T, and E691K) expressed in HEK293T cells reveal distinct effects consistent with the three different biochemical pathways for impairment of AFG3L2 function identified *in vitro*. Non-transfected Mock, wild type (WT), Walker A (WA, ATP binding incompetent), and Walker B (WB, impaired ATP hydrolysis) were included as controls. **C.** ATPase and substrate degradation rates for ^cchex^AFG3L2 and its variants with SCA28-related substitutions in the protease domain show impaired catalytic activity in all cases, albeit to different extents.

### Recessively inherited mutations are involved in substrate cleavage at the proteolytic active site

In contrast to the SCA28 and DOA mutations described above, two single-point mutations in the peptidase domain of AFG3L2 only lead to disease in homozygotes and are linked to severe neurodegenerative disorders with distinct phenotypes (Y616C, SPAX5; A572T, as yet unnamed neurodegenerative disease)^35,36^. Interestingly, we found that both mutations are in close proximity to the peptidase active site (Figure 6A). In AFG3L2, six M41 zinc-metalloprotease active sites are sequestered around the interior periphery of the C6-symmetric proteolytic ring (Figure 6A). At each site a zinc ion is coordinated for catalysis by H574 and H578 from helix α10 and D649 in helix α12 (Figure 6A). The histidine residues are part of a HEXXH motif that is strictly conserved throughout evolution and required for proteolytic activity. Next to this motif is the conserved residue A572, which is directed towards a hydrophobic pocket on the opposite side of α10 from the proteolytic active site (Figures 6A and S5A). The disease mutation A572T would introduce a polar side chain into this hydrophobic pocket, perturbing the local structure of the peptidase active site. In agreement, biochemical analysis of patient mutation ^cchex^AFG3L2 containing the A572T mutations shows significantly decreased protein degradation activity and a moderate defect in ATP hydrolysis (Figure 6B). Moreover, introduction of A572T to ^core^AFG3L2^WB/PI^ did not hamper hexamer recovery by SEC, indicating that A572T causes local structural defects rather than global destabilization (Figure S4E).

**Figure 6.**
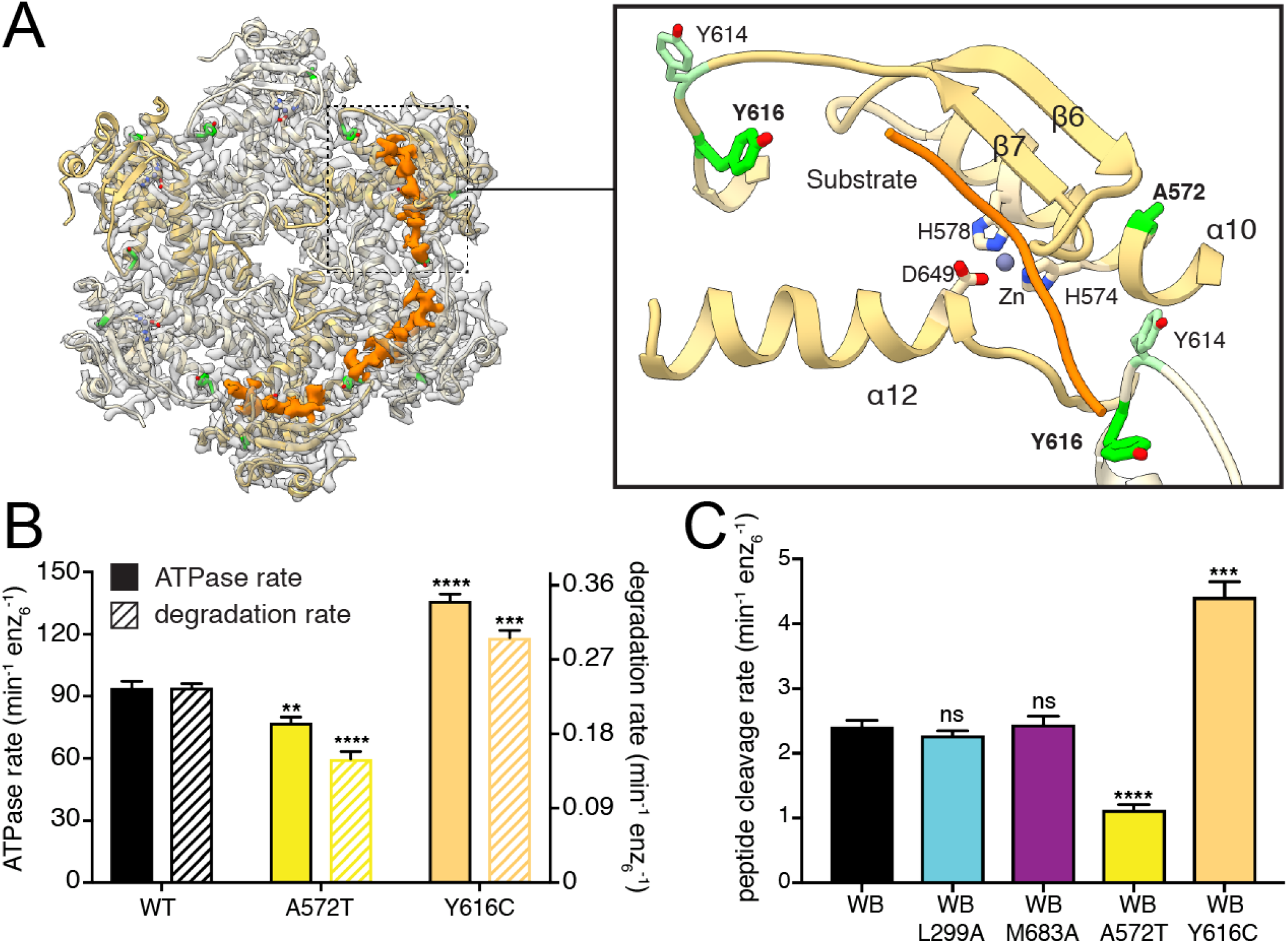
Recessively inherited mutations are directly related to the mechanism of proteolytic cleavage. **A.** Ribbon representation of the atomic model is shown inside the semi-transparent cryo-EM density of the protease ring. Cryo-EM density corresponding to substrate trapped in the proteolytic active sites is displayed as a solid orange isosurface. The close-up view to the right shows how the substrate (orange worm) forms an additional β strand next to β7, and crosses directly above the catalytic triad and the coordinated zinc ion. The disease related residues A572 and Y616 are shown as green sticks, and Y614 is shown in light green. **B.** ATPase and substrate degradation rates for ^cchex^AFG3L2 and its variants containing the Y616C and A572T substitutions related to distinct autosomal recessive conditions show increased and decreased activity, respectively. **C.** Peptide cleavage rate of peptidase active/ATPase inactive AFG3L2 (^core^AFG3L2^WB^) shows that substrate cleavage can occur independently of ATP hydrolysis. Substitutions in the N-termini and the central protease loop (^core^AFG3L2^WB/L299A^ and ^core^AFG3L2^WB/M683A^) do not impact ATP independent proteolytic cleavage, whereas constructs containing the recessive disease mutations A572T and Y616C (^core^AFG3L2^WB/A572T^ and ^core^AFG3L2^WB/Y616C^) present significantly decreased and increased proteolytic activity, respectively.

To confirm that A572T mutation directly affects proteolytic activity of the enzyme, we evaluated peptidase activity independent of ATPase activity using an ATPase-inactive/ peptidase-active variant of the ^core^AFG3L2 protein (denoted ^core^AFG3L2^WB^) to measure the ATPase-independent cleavage of a small fluorogenic peptide as described previously^42^. To validate that this assay is only affected by peptidase activity, we tested the activity of the ^core^AFG3L2^WB/L299A^ and the ^core^AFG3L2^WB/M683A^ constructs. These constructs were selected because we showed that these mutations in the N-terminus and central protrusion, respectively, impair different aspects of substrate processing and decrease substrate degradation rates in the ATPase active construct (Figures 2 and 3). In the ATPase independent assay, neither of these mutations affected the peptide cleavage rate, indicating that this assay reports solely on proteolytic activity independent of upstream mechanistic defects (Figure 6C). As expected, the A572T mutation reduced peptide cleavage rates by 50%, confirming that A572T directly impairs proteolytic cleavage independently of ATPase function (Figure 6C).

Notably, our structure of catalytically inactive AFG3L2 contained density extending across the catalytic triad within three of the six protease active sites, which we attribute to substrate peptide that is positioned for proteolytic cleavage (Figure 6A). This substrate appears to form an additional β-strand next to β7, positioning the targeted polypeptide stably above the catalytic triad. AFG3L2 preferentially cleaves sequences containing phenylalanine residues in the P1’ position, immediately C-terminal of the scissile bond^42^. The position of the bound substrate in our structure locates the putative specificity site (S1’) to a hydrophobic pocket created by V571, L603 and G645 from a single subunit, and L615 from the adjacent subunit (Figure S5B). Although no density is observed for the side chain of the substrate residue closest to this position, a phenylalanine side chain can be easily accommodated in this pocket, consistent with the observed cleavage specificity. Interestingly, L615 is part of a non-conserved loop (Y614, L615 and Y616) at the intersubunit interface, and clear density for the substrate backbone is seen positioned between the side chains of Y614 and the SPAX5-related residue Y616 (Figures 6A and S5B). Strikingly, the disease mutation Y616C displayed significantly increased ATPase and protein degradation rates, with a modest reduction in complex stability (Figure 6B and S4E). Moreover, peptidase activity independent of ATP was two-fold higher for Y616C compared to controls (Figure 6C). These results suggest that substitution of Y616 with a smaller cysteine residue may reduce steric hindrance of incoming polypeptides, thereby increasing accessibility to the S1’ pocket. The distinct gain-of-function effect of the Y616C substitution provides a plausible explanation to the unique phenotype of this mutation in patients.

## Discussion

A hand-over-hand conveyance of substrate along the central pore of a spiraling ATPase ring is rapidly emerging as the conserved mechanism that enables ATP-driven substrate translocation across much of the AAA+ superfamily. However, a fundamental question remains unanswered: “What are the unique evolutionary adaptations that enable different AAA+ proteins to process their distinct substrates?” Our structure of AFG3L2 with substrate bound in both the ATPase and protease domains reveals how an intricate interconnected network within the AFG3L2 hexamer coordinates ATP-driven substrate engagement, translocation, and transfer across the degradation chamber for proteolysis. We show that unique structural features integrate with a core nucleotide-dependent allosteric mechanism to drive substrate processing, and the functional relevance of these distinctive structural features is underscored by our finding that disease-linked mutations are concentrated at these substrate-interacting, non-conserved regions. We further show how disease-relevant substitutions differentially impact AFG3L2 activity and stability, providing a molecular basis for AFG3L2-associated neurodegenerative conditions.

An important first step for AAA+ activity is the recognition of the appropriate substrates for processing. While the C-termini of FtsH-related AAA+ proteases have been shown to modulate substrate specificity, the mechanisms responsible for this modulation are unknown^52^. Our structure reveals that the C-terminus of AFG3L2 establishes interdomain interactions between neighboring subunits that stabilize the hexamer, while simultaneously positioning a highly charged tail for interactions at the membrane surface (Figure 7A). This organization is required for AFG3L2 activity, and is in stark contrast to the organization of the C-terminus in YME1, which we previously observed to be an unstructured loop that would extend away from the protease ring into the IMS (Figure 7B). Thus, we speculate that distinct C-termini evolved in m- and i-AAA proteases, enabling charge-driven interactions for recruitment of membrane-embedded substrates in AFG3L2, and mediating recognition of soluble IMS substrates in YME1. In line with this hypothesis, alteration of the C-terminus of YME1 was previously shown to affect processing of soluble substrates of YME1, but not membrane-embedded substrates^53^. These data provide the structural framework to begin understanding the molecular basis for the differential recruitment of specific substrates by AAA+ proteases.

**Figure 7.**
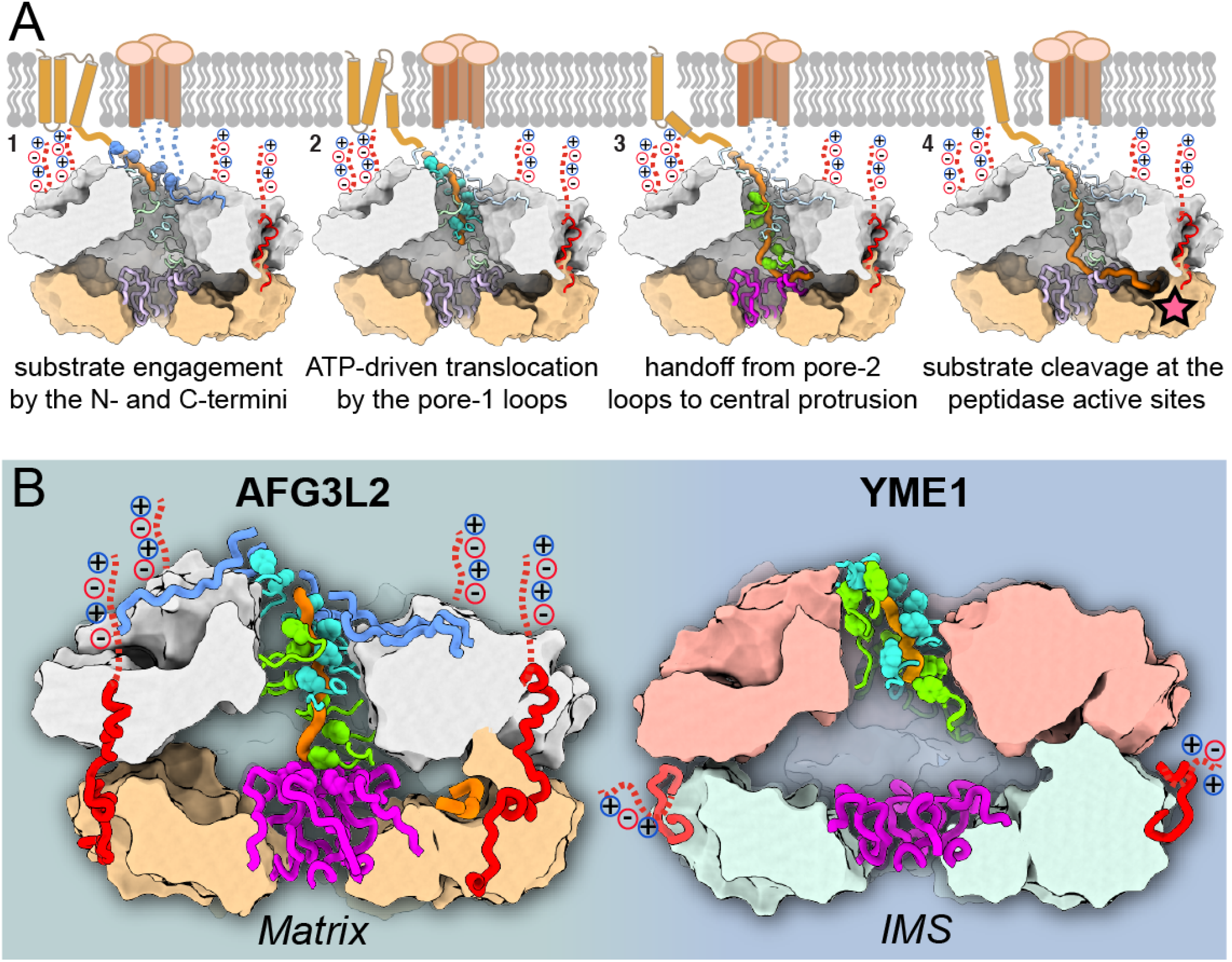
Model for substrate processing by AAA+ proteases of the mitochondrial inner membrane. **A.** Step-by-step model for processing of membrane-associated substrates by AFG3L2. From left to right: (1) The N-and C-termini (in blue and red, respectively) recruit and engage substrate (orange) at the membrane interface; (2) Pore-loops 1 (cyan) intercalate into the substrate to drive hand-over-hand ATP-powered substrate translocation; (3) Pore-loops 2 (green) and the central protrusion of the protease (purple) mediate transfer of the unfolded polypeptide across the proteolytic chamber; (4) The substrate forms an additional β-strand above the zinc coordinated catalytic active site, and is thereby positioned for cleavage (star). **B.** Side-by-side comparison of human m-AAA+ protease AFG3L2 and yeast i-AAA+ protease YME1 emphasizes how small insertions in the pore-loop 2 (green) and central protrusion (purple) dramatically impact the handling of a polypeptide substrate (orange), as well as the organization of the degradation chamber. We further highlight the organization of the non-conserved C-termini (red), and the resulting differential orientation of the charged C-terminal tails (dotted red lines).

In the central pore of the ATPase spiral observed in many AAA+ ATPases, the conserved aromatic residue in pore-loop 1 forms a staircase around the substrate that drives substrate translocation. However, in AFG3L2 the residue immediately adjacent to the pore-loop 1 aromatic links consecutive pore loops to form a tightly packed and continuous chain of residues that surround the substrate. Similarly, both the N-termini of the ATPases and pore-loop 2 are not conserved, and form additional spiral staircases that contact the substrate, mirroring the behavior of pore-loop 1. These three unique structural features of AFG3L2 are integrated with the core machinery for allosteric transmission of ATP-dependent rearrangements, such that all elements can pull on substrate in a concerted fashion in response to ATP hydrolysis (Figure 7A). Additionally, the extended pore-loop 2 of AFG3L2 mediates direct transfer of the substrate to the central protrusion of the protease. Notably, these ancillary substrate-interacting elements are not present in YME1, and, as a result, the translocating substrate enters a large, unoccupied degradation chamber instead (Figure 7B). Thus, our data demonstrate how the cumulative effect of single point substitutions and/or small insertions around the core elements dramatically changes how substrate is handled in different AAA+ proteins (Figure 7B). These changes are likely to affect not only substrate specificity, but also fundamental enzymatic characteristics such as grip on substrate and unfoldase efficacy.

AAA+ proteases are required to degrade a varied collection of substrates with highly diverse sequences. However, evidence exists for site-specific cleavage for some substrates. The presence of substrate in the protease active site in our AFG3L2 structure offers insights into the mechanisms underlying this enzymatic “decision”. We observe that the substrate is stabilized in the active site by predominately backbone-mediated interactions that would be compatible with sequence-independent proteolytic cleavage for broad house-keeping activity. However, our structure also reveals a putative specificity pocket (S1’) that is capable of accommodating an aromatic residue, consistent with the moderate degree of preference for phenylalanine immediately C-terminal of the scissile bond (P1’) reported for AFG3L2^42^. Moreover, the specificity pocket involves residues from the adjacent subunit, suggesting that heterohexameric m-AAA proteases may display altered substrate specificity. These results will guide future studies to determine how distinct substrate interactions in the proteolytic active site might influence sequence preference in different AAA+ proteases, and promote site-specific processing of substrates containing such sequences.

Importantly, how different mutations in AFG3L2 lead to distinct neurological phenotypes is an important, open question in the field. Here, we show that mutations associated with different diseases influence distinct aspects of AFG3L2 stability and activity. Interestingly, the distinct biochemical impact of different mutations appears to correlate with unique phenotypes and inheritance patterns, suggesting that these differences might constitute the molecular basis for the different neurodegenerative conditions associated with AFG3L2 in patients (Table S1). For example, the R468C mutation in DOA patients yields early onset vision impairment without the severe neurological symptoms seen in other AFG3L2-related diseases^33,34^. Incorporation of R468C promotes destabilization of the hexamer both in *vitro* and in vivo, closely mimicking the Walker A mutant that is incapable of binding nucleotide, suggesting that the distinct downstream consequences of impaired nucleotide binding we identify here may be responsible for the DOA phenotype. In contrast, all SCA28-associated mutations in the peptidase domains accumulate close to the central protease protrusion and disrupt intra- or inter-subunit contacts in the proteolytic ring, resulting in destabilization of the hexameric complex and ultimately a broad range of disease severities and onsets that reflect the magnitude of the biochemical defect.

Two recessive mutations found in homozygous patients are positioned close to the proteolytic active site where they produce dramatically different effects on protease activity. Y616C is linked to the rare SPAX5 phenotype and represents the only gain-of-function disease mutation identified for homohexameric AFG3L2 both in our structure-function studies and *in vivo*^35^. Although this mutation produced only a mild destabilizing effect in our AFG3L2 homohexamers, Y616C significantly destabilizes paraplegin-containing heterohexameric m-AAA proteases, consistent with its location close to the intersubunit interface where the mutation could impact the two m-AAA complexes differently^35^. The potential to impact both homo- and heterohexameric m-AAA proteases provides a potential explanation for the overlap of SPAX5 symptoms with those of both SCA28 and hereditary spactic paraplegia (HSP7), which are linked to mutations in AFG3L2 and paraplegin, respectively. Lastly, A572T is the only mutation that appears to directly impair proteolytic cleavage activity and results in an as yet unnamed collection of early on-set and extremely severe neurological symptoms, suggesting that the presence of m-AAA proteases unable to cleave polypeptides at the protease active site have profound consequences in patients^36^. While the functional impact of these different AFG3L2 mutations on mitochondrial proteostasis and function remain to be established, our work provides a molecular basis to begin to understand the distinct phenotypes observed in patients harboring different mutations in this critical AAA+ protease.

## Materials and Methods

### Cloning, Expression, and Purification

Plasmids containing sequences encoding ^cchex^AFG3L2 and ^core^AFG3L2^WB/PI^ were constructed as previously described^42^. These plasmids were used as templates to generate all further mutant variants by site-directed mutagenesis. For biochemical and biophysical characterization assays, ^cchex^AFG3L2, ^core^AFG3L2^WB/PI^, and their variants were expressed in *E. coli* BL21-CodonPlus cells (Agilent) and purified as previously described^42^. For cryo-EM structure determination, ^core^AFG3L2^WB/PI^ was purified as previously described^42^ with the following modifications: Prior to size exclusion chromatography, ^core^AFG3L2^WB/PI^ was exchanged into buffer containing 50 mM HEPES-HCl, pH 7.5, 150 mM KCl, 2 mM EDTA, 10% glycerol and 10 mM β-mercaptoethanol. After buffer exchange, 15 mM MgCl_2_ and 1 mM AMP-PNP were added to the protein sample, and the mixture was incubated for 4 hours on ice. The concentrated protein sample was then applied to Superdex 200 Increase 10/300 GL Increase column (GE Healthcare) equilibrated with buffer containing 25 mM HEPES-HCl, pH 7.5, 100 mM KCl, 20 *μ*M AMP-PNP, 10 mM MgCl_2_, 10% glycerol and 1 mM DTT. Fractions corresponding to the calculated elution volume of ^core^AFG3L2^wb/pi^ hexamer were pooled, concentrated, flash-frozen, and stored at -80 °C. Serving as a model substrate for testing degradation activity of ^cchex^AFG3L2 and its variants,^SF^GFP-10/11^A226G^-β20 was expressed in *E. coli* BL21-CodonPlus cells, and purified as previously described^42^.

### Biochemical Assays

ATPase activity, fluorescence-based protein degradation, and fluorogenic peptide cleavage assays were performed as previously described with some modifications^42,44^. ATPase assays were carried out at 37 °C in a 384-well clear bottom plate (Corning) using a SpectraMax M5 plate reader (Molecular Devices) with 1 *μ*M ^cchex^AFG3L2 or its variants. Steady-state ATPase data were fit to the Hill version of the Michaelis-Menten equation [v = k_ATPase_/(1 + K_0.5_/[ATP]^n^]. Fluorescence based protein degradation assays were performed at 37 ^°^ C in a 384-well black plate (Corning) using a SpectraMax M5 plate reader (ex = 467 nm; em = 511 nm) with 1 *μ*M ^cchex^AFG3L2 or its variants and 20 *μ*M ^SF^GFP-10/11^A226G^-β20. Initial cleavage rates were determined by measuring loss of 511 nm fluorescence over early linear time points. Fluorogenic peptide cleavage assays were performed at 37 ° C with 1 *μ*M ^core^AFG3L2^WB^ or its variants and 50 uM peptide (Leu-(3-NO_2_-Tyr)-Phe-Gly-(Lys-Abz)) (GenScript) in a 384-well black plate using SpectraMax M5 plate reader (ex = 320 nm; em = 420 nm). Initial peptide cleavage rates were calculated from the loss of 420 nm fluorescence over early linear time points.

### Size Exclusion Chromatography (SEC) assay

For determination of SEC profiles, one liter of *E.coli* culture containing ^core^AFG3L2^WB/PI^ or its variants were expressed and purified following the procedure described above for use in biochemical characterization prior to SEC. Proteins were applied to a Superdex 200 10/300 GL Increase column equilibrated with buffer containing 25 mM HEPES-HCl (pH 7.5),100 mM KCl, 10% glycerol, and 1 mM DTT. Elution profiles compared against molecular weight standards to determine the ratio of assembled hexameric species to disassembled species for each ^core^AFG3L2^WB/PI^ variants.

### Sample preparation for electron microscopy

For cryo-EM structure determination, the ^core^AFG3L2^WB/PI^ construct was incubated on ice for 5 minutes in 20mM Tris pH 8, 100mM NaCl, 5mM MgCl_2_, 1mM DTT, 1mM AMPPNP, 1% glycerol and 0.05% Lauryl Maltose Neopentyl Glycol (LMNG, Anatrace). Glycerol and LMNG were added to the buffer to help reduce the preferential orientation of the AFG3L2 particles in vitrified ice. 3 *μ*l of the sample at 2 mg/ml were applied to 300 mesh UltrAuFoil Holey Gold Films R1.2/1.3 (Quantifoil) that had been previously plasma cleaned in a 75% argon / 25% oxygen atmosphere at 15 Watts for 6 seconds using a Solarus plasma cleaner (Gatan, Inc). The grids were loaded into a Vitrobot (ThermoFisher) with an environment chamber at a temperature of 4 ° C and 100% humidity. Grids were blotted for 3 seconds with Whatman No.1 filter paper and plunged into a liquid ethane slurry.

### Electron microscopy data acquisition

Cryo-EM data were collected on a ThermoFisher Talos Arctica transmission electron microscope (TEM) operating at 200keV, which was aligned as previously described^54^. Dose-fractionated movies were collected using a Gatan K2 Summit direct electron detector operating in electron counting mode, saving 30 frames during a 7s exposure. At an exposure rate of 4.3 e^−^/pixel, this resulted in a cumulative exposure of 40 e^−^/Å^2^. The Leginon data collection software^55^ was used to collect 5,707 micrographs by moving to the center of four holes and image shifting to acquire four exposures at 36,000x nominal magnification (1.15 Å/pixel at the specimen level), with a nominal defocus range of -0.8 to -1.8 *μ*m.

### Image processing

The Appion image processing workflow^56^ was used to perform real-time image pre-processing during cryo-EM data collection. Micrograph frames were aligned using MotionCorr2^57^ and CTF parameters were estimated with CTFFind4^58^. Only micrographs with Appion confidence values above 95% were further processed. A small subset of ~20,000 particles were selected with the FindEM template-based particle picker^59^, using the negative stain 2D classes as templates. This subset of particles was extracted using an unbinned box size of 256 pixels, and binned by a factor of four for 2D classification using RELION 1.4^60^. The resulting class averages were used as templates to select particles using FindEM, yielding 4,541,491 coordinates for putative particle.

Relion 3.0^61^ was used for all the remaining processing steps (Figure S1B). Particles were extracted using an unbinned box size of 160 pixels and binned by a factor of four for 2D classification. The resulting class averages showed a preferred hexameric view of the complex, so only the particles contributing to one of these hexameric classes were retained for subsequent processing steps. All other particles contributing to 2D classes containing well-defined secondary structural elements were also selected for 3D analysis. A total of 2,901,805 particles were extracted (binned by two, box size 80 pixels) and classified into five classes using the scaled and low-pass filtered cryo-EM structure of yeast YME1 (EMDB-7023^8^ as an initial model and a regularization parameter of eight. The 1,129,437 particles contributing to the three best 3D class averages containing well-defined structural features were extracted with a box size of 240 pixels using centered coordinates. These particles were refined to a resolution of ~3.2 Å resolution, although the observable structural features were not consistent with this reported resolution, and were not sufficiently resolved for atomic modeling.

We next performed CTF refinement and beam tilt estimation (estimated beam tilt x=-0.25, beam tilt y=0.07), and used these estimates to re-refine the particle stack, which yielded a structure with a reported resolution of ~3.0 Å. However, this reconstruction still appeared to suffer from issues with flexibility and misalignment, and did not contain structural features that are consistent with 3.0 Å cryo-EM density. Suspecting that these apparent misalignments may be due to subtle motions of the ATPase and protease rings relative to one another, we next performed a multi-body refinement of the AFG3L2 hexamer using three masked regions: the c6-symmetric protease ring, the four most stable ATPase domains (subunits B-E), and the two remaining ATPase domains for which the cryo-EM density was poorly resolved (subunits A,F). While these two flexible domains were not improved by the multibody refinement approach and are reported at ~5 Å resolution, the observable details of the protease and four stable ATPase domains improved substantially. The reported resolutions for the ATPase and protease domains are ~3.0 Å and ~2.9 Å, respectively, by FSC 0.143.

Local resolution estimation using the “blocres” function in the Bsoft package^62^ indicates that there are regions within the reconstructions that are resolved to better than the FSC-reported resolutions. However, we do not observe EM density corresponding to ordered water molecules, which should be visible at better than 2.9 Å resolution so we suspect that the reported FSC-based resolutions are slightly inflated, possibly due to the presence of preferred views in the final reconstruction (Figure S1D). The individual reconstructions output from the multi-body refinement were individually sharpened and low-pass filtered (Table S2), and a composite map was generated to facilitate atomic model building using the “vop max” function in UCSF Chimera^63^.

### Atomic model building and refinement

A homology model was generated using the structure of a subunit of YME1 (PDB ID: 6AZ0) as a starting point. Six copies of this atomic model were generated and each split into the three domains and rigid body fit into each subunit using the “fit in map” function in UCSF Chimera^63^. The ATPase domains of the apo and ATP1 subunits (subunits F and A, respectively) were not further processed as the quality of the density for these domains did not allow for confident atomic modeling, and are included in the deposited model only as a Cα trace. The homology model for the rest of the hexamer, including nucleotides and coordinating metals, was adjusted using the COOT software package^64^, and then real-space refined with rigid body fitting and simulated annealing using the PHENIX package^65^. This refined model was used as input for a multi-model-generating pipeline^66^ that aided in assessing the quality of the model and the map for mechanistic interpretation (Figure S1F). Briefly, the refined model and the map were used to generate 200 models in Rosetta, and the top ten scoring models were selected for further refinement in Phenix. The per-residue Cα root mean square deviation (RMSD) of top ten models was calculated and used to identify regions with poor model convergence. These regions were inspected and re-modeled and re-refined. Regions of the map with poor convergence either had the side-chains were truncated to the Cβ or removed entirely from the model. UCSF Chimera^63^ was used to generate the figures.

### Cell culture and mitochondrial isolation

HEK293T cells were cultured in DMEM (Cellgro) supplemented with 10% fetal bovine serum (Cellgro), 1% penicillin/streptavidin (Gibco) and 1% L-glutamine (Cellgro) at 37 °C (5% CO_2_). Cells were transiently transfected with a pcDNA3.1 vector that expressed a C-terminally Flag-tagged AFG3L2 (GenScript) with the indicated single-point mutations. After 36h, mitochondria were isolated from cells as previously reported^67^.

### SDS- and BN-PAGE analysis

Freshly isolated mitochondria were lysed in SDS (20 mM Hepes pH 7.5, 100 mM NaCl, 1 mM EDTA, 1% Triton supplemented with EDTA-free protease inhibitors (Roche)) or BN-Lysis buffer (20 mM Tris pH 7.5, 10% w/v glycerol, 50 mM NaCl, 0.1 mM EDTA, and 1mM PMSF supplemented with 1% dodecyl-maltoside). Samples were normalized by total protein concentration using Bio-Rad protein quantification. For SDS-PAGE, lysates were separated on Tris-glycine gels and transferred onto nitrocellulose membranes (Bio-Rad) for immunoblotting. Following incubation with primary antibodies, membranes were incubated with IR-labeled secondary antibodies (Li-COR Biosciences) and analyzed using the Odyssey Infrared Imaging System (Li-COR Biosciences). BN-PAGE samples were separated on 4-16% polyacrylamide gels, and western blotted as described^68^. Primary antibodies: anti-AFG3L2 (Genetex, cat. GTX102036, 1:1000 dilution), anti-Flag (M2 Flag, Sigma, cat. F18094, 1:1000 dilution), anti-Tubulin (Sigma, cat. T6074, 1:10000 dilution).

## Acknowledgements

We thank J.C. Ducom at The Scripps Research Institute High Performance Computing for computational support, and B. Anderson for electron microscope support. C.P. is supported by an American Heart Association predoctoral fellowship. We thank M. Shin for helpful discussion. G.C.L. is supported as a Pew Scholar, by an Amgen Young Investigator award, and the National Institutes of Health (NIH) DP2EB020402. Computational analyses of EM data were performed using shared instrumentation funded by NIH S10OD021634 to G.C.L. S.E.G. is supported by NIH R01GM115898. R.L.W. is supported by NIH R01NS095892.

## Data Availability

All the cryo-EM maps were deposited in the Electron Microscopy Data Bank under accession code EMD-0552. The associated atomic coordinates were deposited into the Protein Data Bank with accession code 6NYY.

## Supplementary Materials

**Figure S1.**
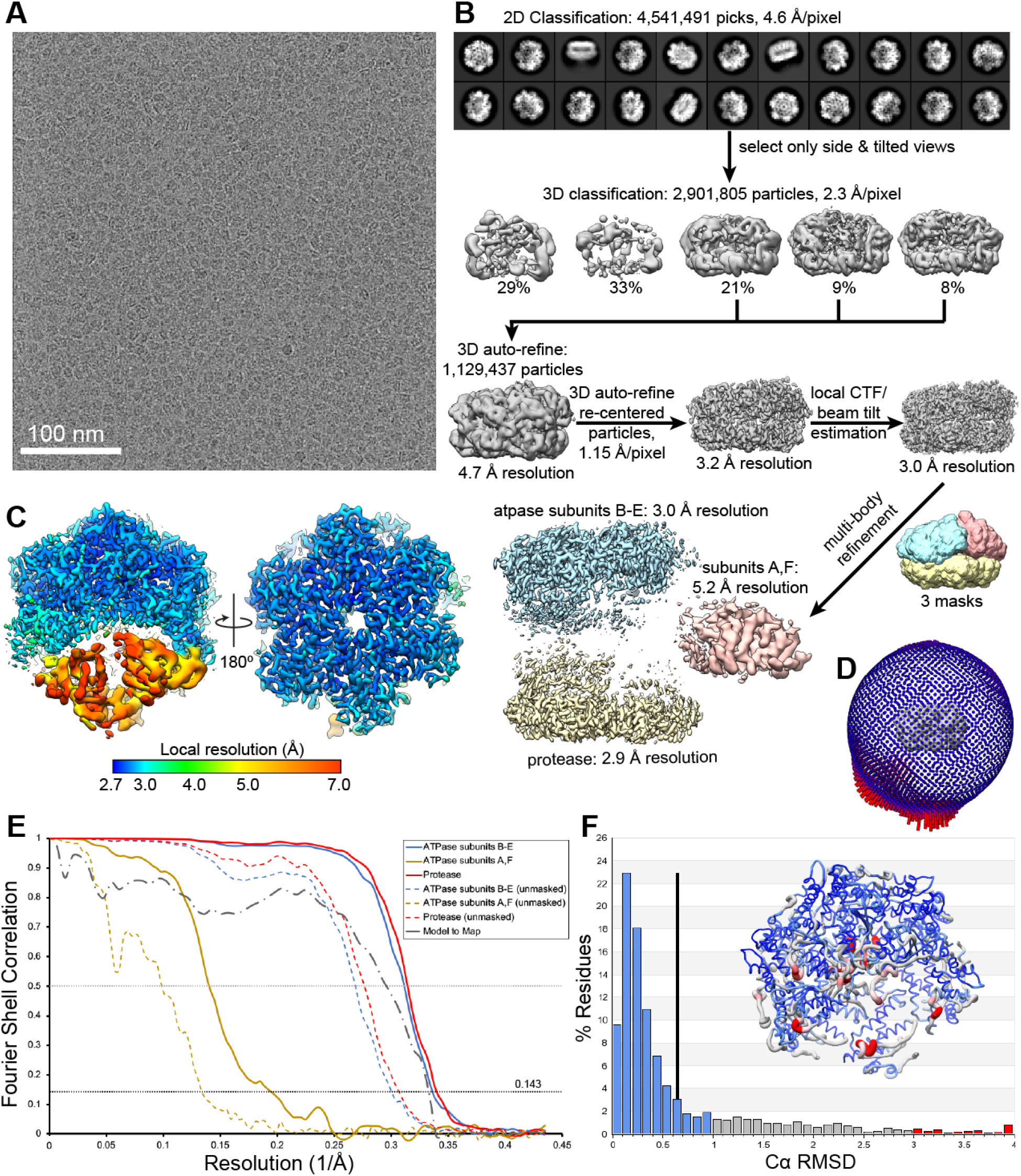
Structure determination and validation. **A.** Representative motion-corrected, dose-weighted cryo-EM micrograph of vitrified substrate-bound ^core^AFG3L2^WB/PI^. **B.** RELION processing scheme used to determine the 3D reconstruction of AFG3L2. Details provided in the Methods. **C.** Local resolution were calculated using the Bsoft package^62^, showing that the final EM-density of ^core^ AFG3L2^WB/PI^ shows the resolution varies from 3.1 Å at the core of the complex to 3.5 Å for the built regions. **D.** The distribution of the Euler angles assigned to the particles in the final consensus reconstruction. Note the presence of some preferred views, which may inflate the reported FSC-based resolution. **E.** FSC plots of the unmasked and masked reconstructions of the three components refined through multibody analysis in RELION. The FSC calculated using the composite map and the atomic model is also shown (dashed grey line). **F.** Histogram of the per-residue Cα RMSD values calculated from the top ten refined atomic models with the mean per-residue Cα RMSD value shown as a black vertical bar^66^. Inset is a worm representation of the atomic model, colored according to the per-residue Cα RMSD values depicted in the histogram.

**Figure S2.**
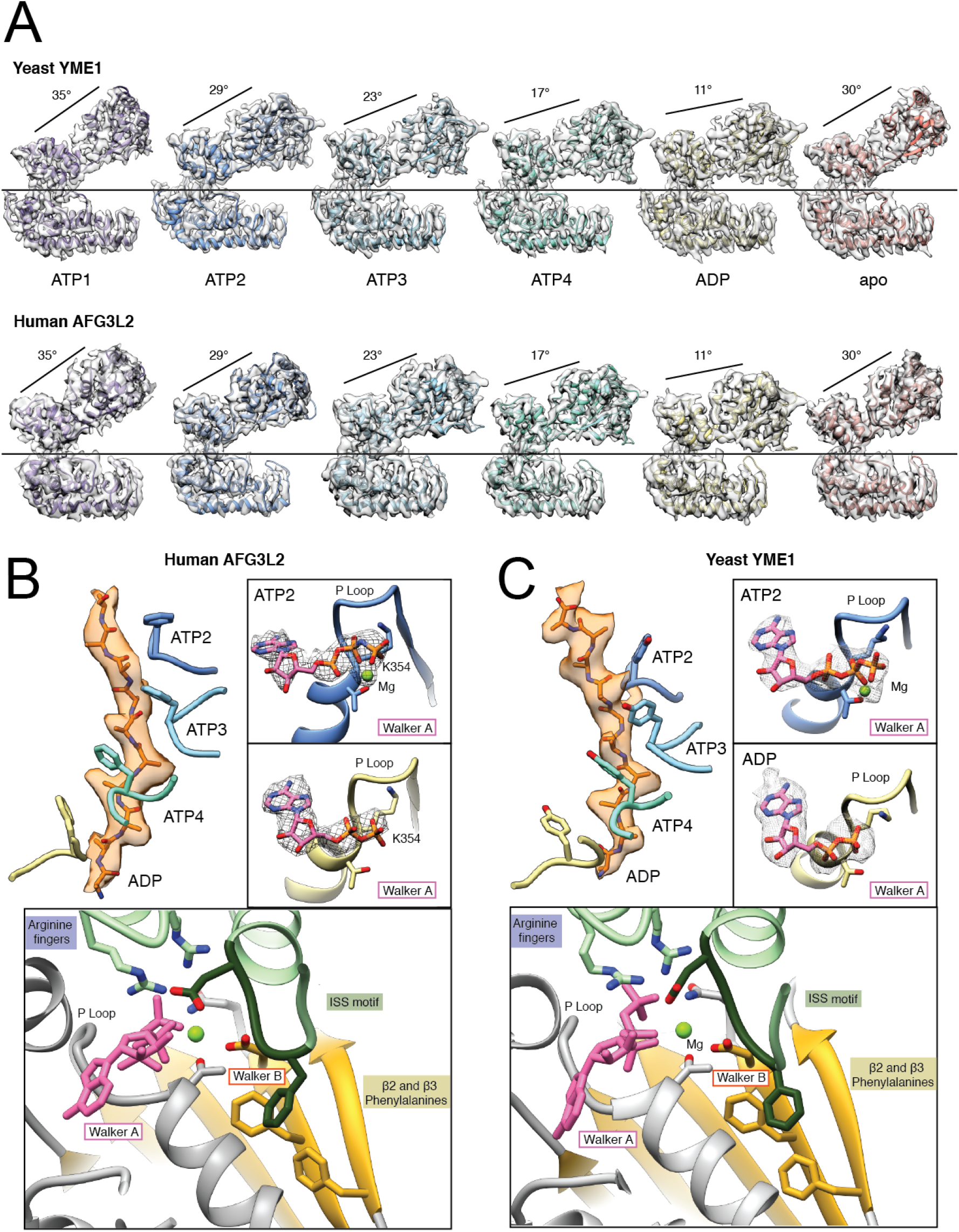
Conservation of the hand-over-hand mechanism for substrate translocation across eukaryotes. **A.** Individual protomers of yeast i-AAA protease YME1 (top) and human m-AAA protease AFG3L2 (bottom) shown side by side, aligned with the protease domain in the same orientation. The cryo-EM density is shown as a transparent grey isosurface. For each subunit, the relative orientation between the ATPase and protease domains is almost identical in YME1 and AFG3L2. The core mechanistic features of **B.** AFG3L2 and **C.** YME1 are essentially indistinguishable. Top left, the conserved pore-loop 1 aromatic residue intercalates into the substrate backbone every two amino acids. Top right, the nucleotide binding features characteristic of P-loop ATPases are shown for the ATP2 and ADP subunits with the cryo-EM density corresponding to nucleotide shown in mesh. Bottom, the ATP-bound nucleotide-binding pocket reveals conservation of all the elements involved in the ATP hydrolysis cycle and the allosteric transmission of the nucleotide-dependent conformational changes to the pore-loops.

**Figure S3.**
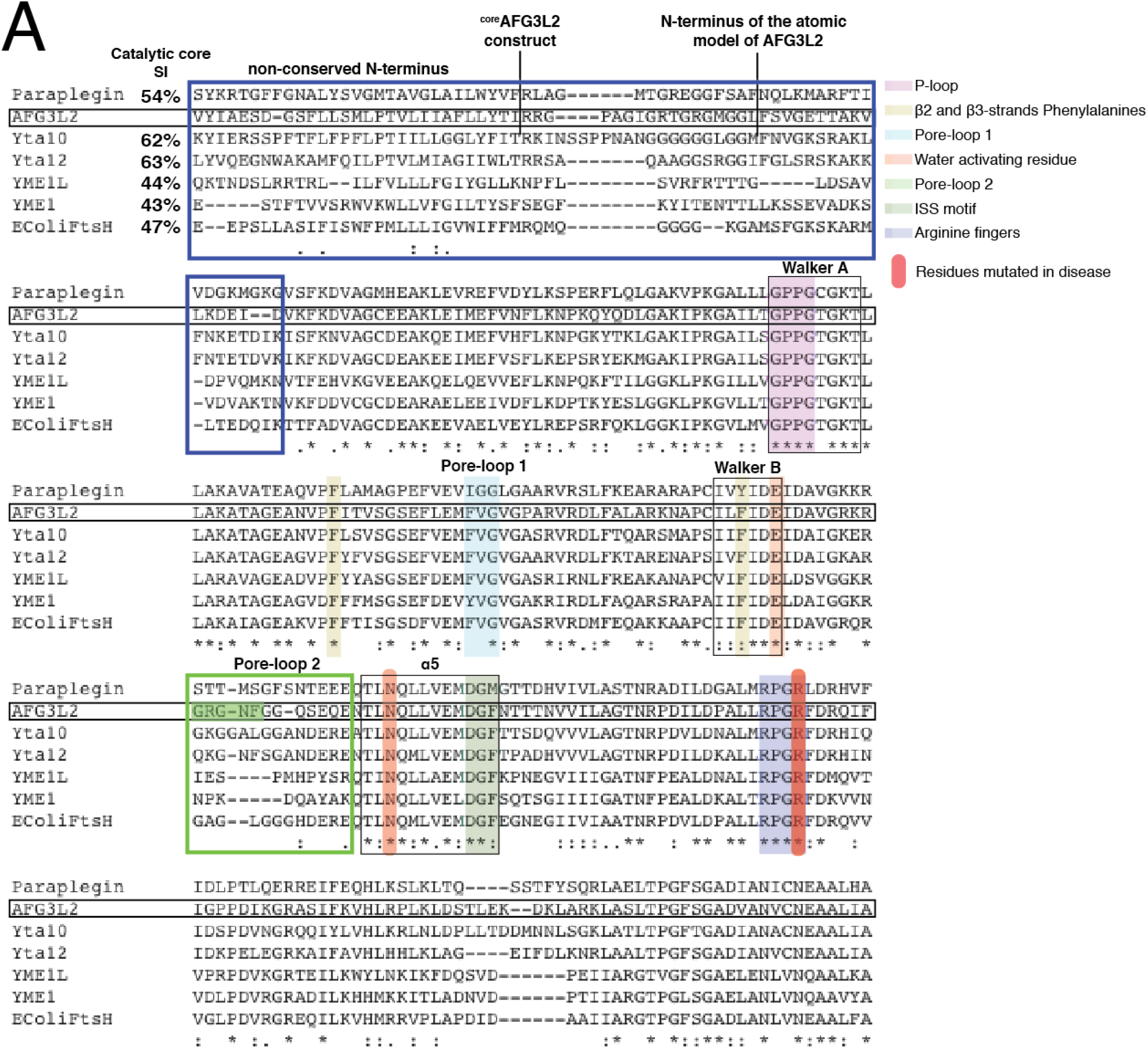
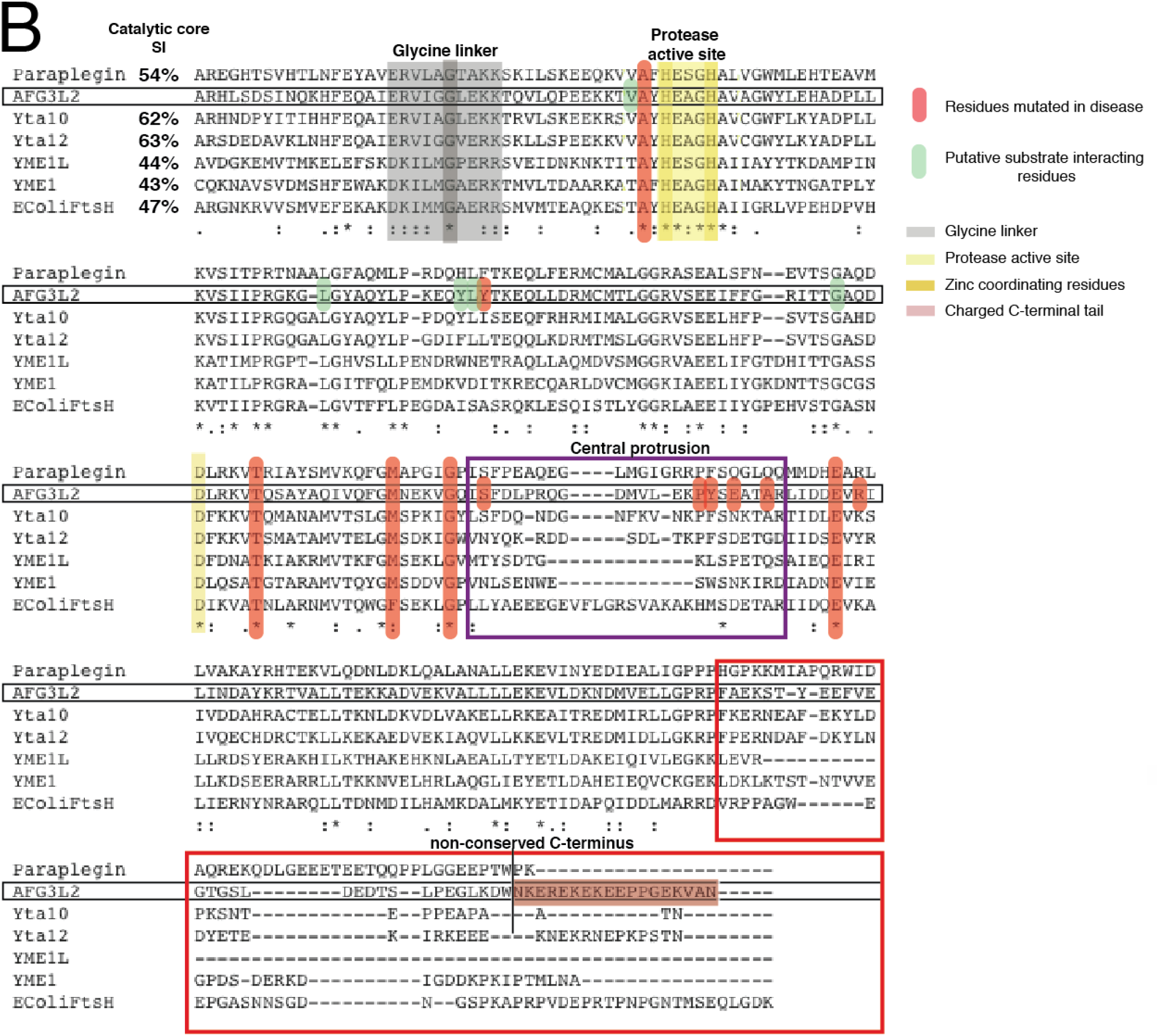
Sequence analysis of the FtsH-related protein family. ClustalW alignment of the Uniprot sequences of **A.** the ATPase and **B.** the protease domains of human AFG3L2 and diverse members of FtsH-related proteins: human paraplegin, *S. cerevisiae* m-AAA proteases Yta10 and Yta12, human and *S. cerevisiae* i-AAA proteases YME1L and YME1, and *E. coli* FtsH. We specify overall sequence identity of the catalytic core of each homolog to AFG3L2 (in %) and, under each residue, an asterisk denotes strict conservation across all homologs, and one or two dots indicate high sequence similarity. All elements identified to be important for the mechanism of ATP hydrolysis, substrate translocation, and proteolytic cleavage are strictly conserved and labeled in the figure legend. Disease-related residues in AFG3L2 are highlighted in red and the non-conserved regions of the catalytic core are delineated by boxes, colored as follows: N-terminus (blue), pore-loop 2 (green), central protease loop (purple), and C-terminus (red).

**Figure S4.**
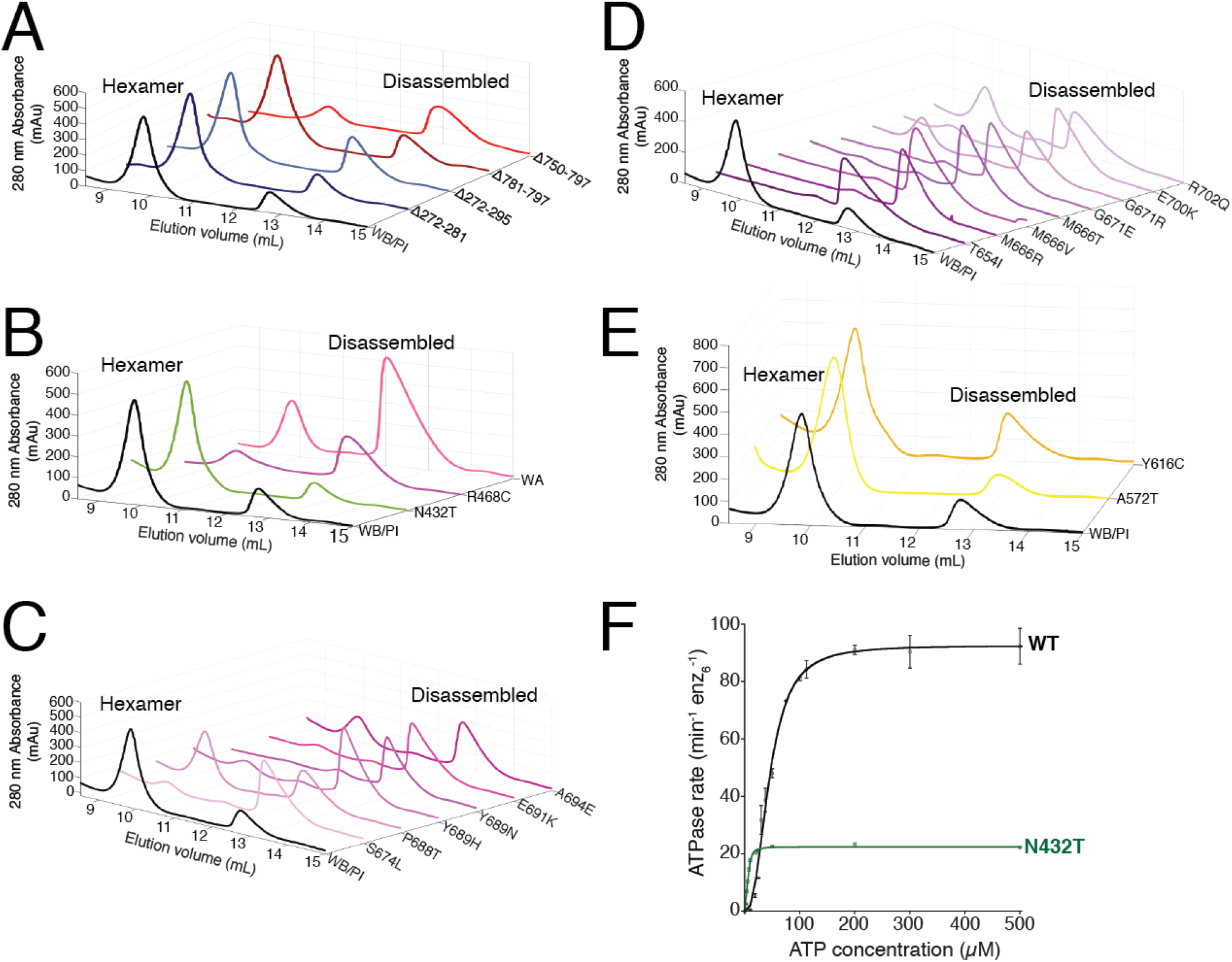
The effects of diverse mutations on AFG3L2 hexameric stability and ATP binding/hydrolysis *in vitro*. Size exclusion chromatography (SEC) analysis of the ^core^AFG3L2^WB/PI^ construct upon incorporation of **A.** Truncations in the N- and C-termini, **B.** the ATPase disease-related substitutions with the ATP binding incompetent Walker A (WA, K354A) as a control, C. the mutations of the central protrusion associated with disease, D. the disease-related substitutions of the lateral interface in between adjacent protease domains, and E. the recessive disease-linked substitutions involved in proteolytic cleavage. In each case, the ^core^AFG3L2^WB/PI^ construct was used as a control and the loss of the hexameric peak and increase in the disassembled peak indicate reduced stability of the affected constructs. F. Rate of ATP hydrolysis against ATP concentration for ^cchex^AFG3L2 and its variant with N432T substitution. Data were fit to the Hill version of the Michaelis-Menten equation (For WT: *k*_ATPase_ = 92.6 min^−1^ enz_6_^−1^; *K*_0.5_ = 45 *μ*M; For N432T: *k*_ATPase_ = 22.4 min^−1^ enz_6_^−1^; *K*_0.5_ = 5.6 *μ*M).

**Figure S5.**
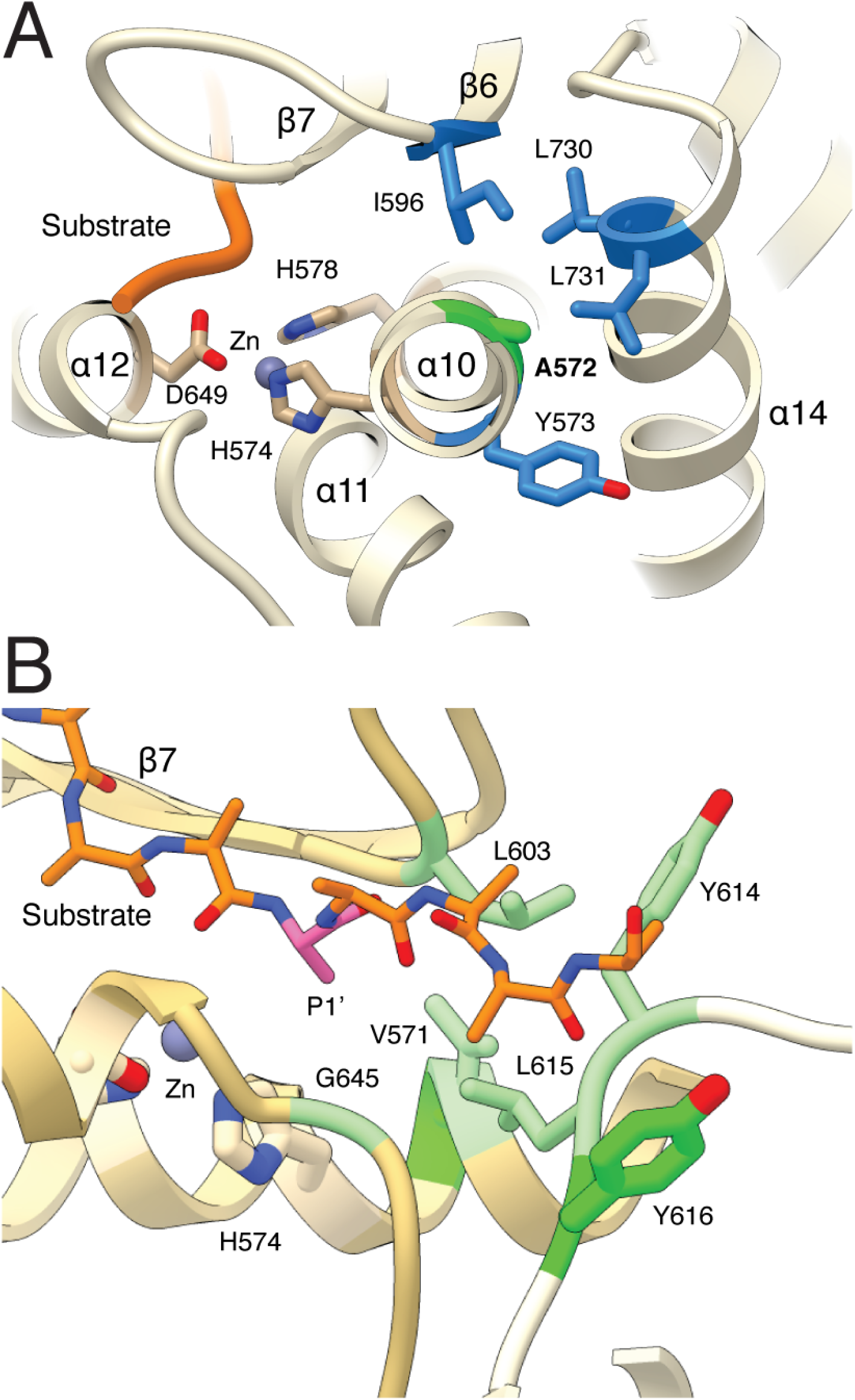
Structural details of the protease active site. **A.** Side view of the proteolytic site shows that the disease-related residue A572 (highlighted in green) inserts into a hydrophobic pocket formed by residues Y573, I596, L730 and L731, all of which are shown in blue. **B.** Top view of the proteolytic site shows that the P1’ residue (pink) of the substrate faces a hydrophobic pocket (green) formed by residues V571, L603, G645 of the *cis* subunit, and L615 from the adjacent subunit. The substrate density extends in between Y614 and the disease-associated Y616 from the neighboring subunit.

**Table S1.**
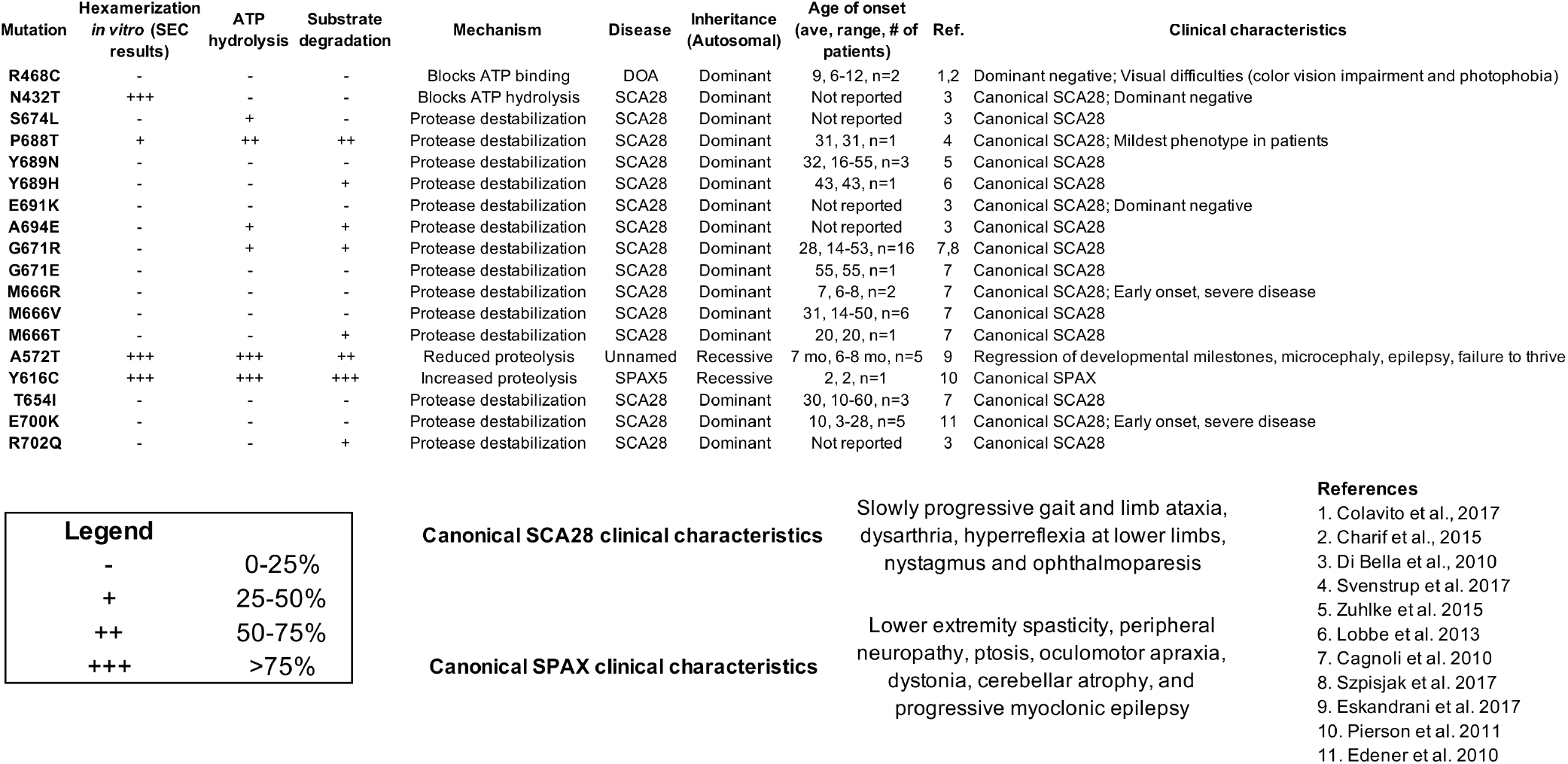
Biochemical characterization and reported clinical phenotypes linked to disease-related substitutions in AFG3L2. For each mutation characterized in this study, we show the hexamerization, ATP hydrolysis and substrate degradation rates we determined in vitro, as well as our proposed mechanism. We also summarize and reference the clinical characteristics and inheritance patterns previously reported in each case.

Unique structural features of the mitochondrial AAA+ protease AFG3L2 reveal the molecular basis for activity in health and disease
| Mutation | Hexamerization<br>*in vitro* (SEC<br>results) | ATP<br>hydrolysis | Substrate<br>degradation | Mechanism | Disease | Inheritance<br>(Autosomal) | Age of onset<br>(ave, range, # of<br>patients) | Ref. | Clinical characteristics |
| --- | --- | --- | --- | --- | --- | --- | --- | --- | --- |
| R468C | - | - | - | Blocks ATP binding | DOA | Dominant | 9, 6-12, n=2 | 1,2 | Dominant negative; Visual difficulties (color vision impairment and photophobia) |
| N432T | +++ | - | - | Blocks ATP hydrolysis | SCA28 | Dominant | Not reported | 3 | Canonical SCA28; Dominant negative |
| S674L | - | + | - | Protease destabilization | SCA28 | Dominant | Not reported | 3 | Canonical SCA28 |
| P688T | + | ++ | ++ | Protease destabilization | SCA28 | Dominant | 31, 31, n=1 | 4 | Canonical SCA28; Mildest phenotype in patients |
| Y689N | - | - | - | Protease destabilization | SCA28 | Dominant | 32, 16-55, n=3 | 5 | Canonical SCA28 |
| Y689H | - | - | + | Protease destabilization | SCA28 | Dominant | 43, 43, n=1 | 6 | Canonical SCA28 |
| E691K | - | - | - | Protease destabilization | SCA28 | Dominant | Not reported | 3 | Canonical SCA28; Dominant negative |
| A694E | - | + | + | Protease destabilization | SCA28 | Dominant | Not reported | 3 | Canonical SCA28 |
| G671R | - | + | + | Protease destabilization | SCA28 | Dominant | 28, 14-53, n=16 | 7,8 | Canonical SCA28 |
| G671E | - | - | - | Protease destabilization | SCA28 | Dominant | 55, 55, n=1 | 7 | Canonical SCA28 |
| M666R | - | - | - | Protease destabilization | SCA28 | Dominant | 7, 6-8, n=2 | 7 | Canonical SCA28; Early onset, severe disease |
| M666V | - | - | - | Protease destabilization | SCA28 | Dominant | 31, 14-50, n=6 | 7 | Canonical SCA28 |
| M666T | - | - | + | Protease destabilization | SCA28 | Dominant | 20, 20, n=1 | 7 | Canonical SCA28 |
| A572T | +++ | +++ | ++ | Reduced proteolysis | Unnamed | Recessive | 7 mo, 6-8 mo, n=5 | 9 | Regression of developmental milestones, microcephaly, epilepsy, failure to thrive |
| Y616C | +++ | +++ | +++ | Increased proteolysis | SPAX5 | Recessive | 2, 2, n=1 | 10 | Canonical SPAX |
| T654I | - | - | - | Protease destabilization | SCA28 | Dominant | 30, 10-60, n=3 | 7 | Canonical SCA28 |
| E700K | - | - | - | Protease destabilization | SCA28 | Dominant | 10, 3-28, n=5 | 11 | Canonical SCA28; Early onset, severe disease |
| R702Q | - | - | + | Protease destabilization | SCA28 | Dominant | Not reported | 3 | Canonical SCA28 |

Legend
|  |  |
| --- | --- |
| - | 0-25% |
| + | 25-50% |
| ++ | 50-75% |
| +++ | >75% |

**Table S2.** Cryo-EM data collection, refinement, and validation statistics

|  |  |  |  |
| --- | --- | --- | --- |
| core AFG3L2<sup>WB/PI</sup> |  |  |  |
| PDB ID: 6NYY |  |  |  |
| EMDB ID: 0552 |  |  |  |
| Data Collection |  |  |  |
| Microscope | Talos Arctica |  |  |
| Camera | K2 Summit |  |  |
| Magnification | 36,000 |  |  |
| Voltage (kV) | 200 |  |  |
| Total electron exposure (e<sup>-</sup>/Å<sup>2</sup>) | 40 |  |  |
| Exposure rate (e<sup>-</sup>/pixel/sec) | 4.3 |  |  |
| Defocus range (μm) | 0.8 to 1.8 |  |  |
| Pixel size (Å/pixel)* | 1.15 |  |  |
| Micrographs collected (no.) | 5,707 |  |  |
| Micrographs used (no.) | 5,378 |  |  |
| Total extracted picks (no.) | 4,541,491 |  |  |
| Refined particles (no.) | 2,901,805 |  |  |
| Reconstruction |  |  |  |
| Final Particles (no.) | 1,129,437 |  |  |
| Symmetry | C1 |  |  |
| Resolution (global) (Å) |  |  |  |
| FSC 0.5 (unmasked/masked) | ATPase: 3.7/3.2 | ATPase A,F: 10.1/7.1 | Protease: 3.6/3.2 |
| FSC 0.143 (unmasked/masked) | ATPase B-E: 3.2/3.0 | ATPase A,F: 7.5/5.2 | Protease: 3.3/2.9 |
| Local resolution range (Å) | 2.7 - 10 |  |  |
| Map sharpening *B*-factor (Å<sup>2</sup>) | ATPase B-E: -111 | ATPase A,F: -155 | Protease: -93 |
| Model composition* |  |  |  |
| Nonhydrogen atoms | 18,704 |  |  |
| Protein residues | 2,396 |  |  |
| Ligands | 12 |  |  |
| Refinement |  |  |  |
| MapCC (global/local) | 0.84/0.85 |  |  |
| *B*-factors (Å<sup>2</sup>) |  |  |  |
| Nonhydrogen atoms | 84.6 |  |  |
| Protein residues | 84.6 |  |  |
| Ligands | 75.3 |  |  |
| R.m.s. deviations |  |  |  |
| Bond lengths (Å) | 0.006 |  |  |
| Bond angles (°) | 0.799 |  |  |
| Validation |  |  |  |
| MolProbity score | 1.58 |  |  |
| Clashscore | 3.93 |  |  |
| Poor rotamers (%) | 0.05 |  |  |
| Ramachandran plot |  |  |  |
| Favored (%) | 94.06 |  |  |
| Allowed (%) | 5.94 |  |  |
| Disallowed (%) | 0.00 |  |  |
| EMRinger<sup>66</sup> score | 3.2 |  |  |

\*Atomic statistics and validation don't include the ATPase domains of subunits A & F

## References

1. Chen, B., Retzlaff, M., Roos, T. & Frydman, J. Cellular strategies of protein quality control. Cold Spring Harb Perspect Biol 3, a004374, doi:10.1101/cshperspect.a004374 (2011).

2. Bohovych, I., Chan, S. S. & Khalimonchuk, O. Mitochondrial protein quality control: the mechanisms guarding mitochondrial health. Antioxid Redox Signal 22, 977–994, doi:10.1089/ars.2014.6199 (2015).

3. Langklotz, S., Baumann, U. & Narberhaus, F. Structure and function of the bacterial AAA protease FtsH. Biochim Biophys Acta 1823, 40–48, doi:10.1016/j.bbamcr.2011.08.015 (2012).

4. Leonhard, K. et al. AAA proteases with catalytic sites on opposite membrane surfaces comprise a proteolytic system for the ATP-dependent degradation of inner membrane proteins in mitochondria. EMBO J 15, 4218–4229 (1996).

5. Wagner, R., Aigner, H. & Funk, C. FtsH proteases located in the plant chloroplast. Physiol Plant 145, 203–214, doi:10.1111/j.1399-3054.2011.01548.x (2012).

6. Sauer, R. T. & Baker, T. A. AAA+ proteases: ATP-fueled machines of protein destruction. Annu Rev Biochem 80, 587–612, doi:10.1146/annurev-biochem-060408-172623 (2011).

7. Janska, H., Kwasniak, M. & Szczepanowska, J. Protein quality control in organelles - AAA/FtsH story. Biochim Biophys Acta 1833, 381–387, doi:10.1016/j.bbamcr.2012.03.016 (2013).

8. Puchades, C. et al. Structure of the mitochondrial inner membrane AAA+ protease YME1 gives insight into substrate processing. Science 358, doi:10.1126/science.aao0464 (2017).

9. Bittner, L. M., Arends, J. & Narberhaus, F. When, how and why? Regulated proteolysis by the essential FtsH protease in Escherichia coli. Biol Chem 398, 625–635, doi:10.1515/hsz-2016-0302 (2017).

10. Glynn, S. E. Multifunctional Mitochondrial AAA Proteases. Front Mol Biosci 4, 34, doi:10.3389/fmolb.2017.00034 (2017).

11. Nishimura, K., Kato, Y. & Sakamoto, W. Chloroplast Proteases: Updates on Proteolysis within and across Suborganellar Compartments. Plant Physiology 171, 2280–2293, doi:10.1104/pp.16.00330 (2016).

12. Banfi, S. et al. Identification and characterization of AFG3L2, a novel paraplegin-related gene. Genomics 59, 51–58, doi:10.1006/geno.1999.5818 (1999).

13. Koppen, M., Metodiev, M. D., Casari, G., Rugarli, E. I. & Langer, T. Variable and tissue-specific subunit composition of mitochondrial m-AAA protease complexes linked to hereditary spastic paraplegia. Mol Cell Biol 27, 758–767, doi:10.1128/MCB.01470-06 (2007).

14. Nolden, M. et al. The m-AAA protease defective in hereditary spastic paraplegia controls ribosome assembly in mitochondria. Cell 123, 277–289, doi:10.1016/j.cell.2005.08.003 (2005).

15. Arlt, H. et al. The formation of respiratory chain complexes in mitochondria is under the proteolytic control of the m-AAA protease. Embo Journal 17, 4837–4847, doi:DOI 10.1093/emboj/17.16.4837 (1998).

16. Consolato, F., Maltecca, F., Tulli, S., Sambri, I. & Casari, G. m-AAA and i-AAA complexes coordinate to regulate OMA1, the stress-activated supervisor of mitochondrial dynamics. Journal of Cell Science 131, doi:UNSP jcs213546 10.1242/jcs.213546 (2018).

17. Konig, T. et al. The m-AAA Protease Associated with Neurodegeneration Limits MCU Activity in Mitochondria. Molecular Cell 64, 148–162, doi:10.1016/j.molcel.2016.08.020 (2016).

18. Maltecca, F. et al. The mitochondrial protease AFG3L2 is essential for axonal development. Journal of Neuroscience 28, 2827–2836, doi:10.1523/Jneurosci.4677-07.2008 (2008).

19. Maltecca, F. et al. Haploinsufficiency of AFG3L2, the Gene Responsible for Spinocerebellar Ataxia Type 28, Causes Mitochondria-Mediated Purkinje Cell Dark Degeneration. Journal of Neuroscience 29, 9244–9254, doi:10.1523/Jneurosci.1532-09.2009 (2009).

20. Almajan, E. R. et al. AFG3L2 supports mitochondrial protein synthesis and Purkinje cell survival. Journal of Clinical Investigation 122, 4048–4058, doi:10.1172/Jci64604 (2012).

21. Kondadi, A. K. et al. Loss of the m-AAA protease subunit AFG3L2 causes mitochondrial transport defects and tau hyperphosphorylation. Embo Journal 33, 1011–1026, doi:10.1002/embj.201387009 (2014).

22. Mancini, C. et al. Mice harbouring a SCA28 patient mutation in AFG3L2 develop late-onset ataxia associated with enhanced mitochondrial proteotoxicity. Neurobiol Dis 124, 14–28, doi:10.1016/j.nbd.2018.10.018 (2018).

23. Levytskyy, R. M., Germany, E. M. & Khalimonchuk, O. Mitochondrial Quality Control Proteases in Neuronal Welfare. J Neuroimmune Pharmacol 11, 629–644, doi:10.1007/s11481-016-9683-8 (2016).

24. Mariotti, C. et al. Spinocerebellar ataxia type 28: a novel autosomal dominant cerebellar ataxia characterized by slow progression and ophthalmoparesis. Cerebellum 7, 184–188, doi:10.1007/s12311-008-0053-9 (2008).

25. Di Bella, D. et al. Mutations in the mitochondrial protease gene AFG3L2 cause dominant hereditary ataxia SCA28. Nat Genet 42, 313–321, doi:10.1038/ng.544 (2010).

26. Cagnoli, C. et al. Missense Mutations in the AFG3L2 Proteolytic Domain Account for similar to 1.5% of European Autosomal Dominant Cerebellar Ataxias. Human Mutation 31, 1117–1124, doi:10.1002/humu.21342 (2010).

27. Lobbe, A. M. et al. A Novel Missense Mutation in AFG3L2 Associated with Late Onset and Slow Progression of Spinocerebellar Ataxia Type 28. Journal of Molecular Neuroscience 52, 493–496, doi:10.1007/s12031-013-0187-1 (2014).

28. Zuhlke, C. et al. Spinocerebellar ataxia 28: a novel AFG3L2 mutation in a German family with young onset, slow progression and saccadic slowing. Cerebellum Ataxias 2, 19, doi:10.1186/s40673-015-0038-7 (2015).

29. Svenstrup, K. et al. SCA28: Novel Mutation in the AFG3L2 Proteolytic Domain Causes a Mild Cerebellar Syndrome with Selective Type-1 Muscle Fiber Atrophy. Cerebellum 16, 62–67, doi:10.1007/s12311-016-0765-1 (2017).

30. Szpisjak, L. et al. Neurocognitive Characterization of an SCA28 Family Caused by a Novel AFG3L2 Gene Mutation. Cerebellum 16, 979–985, doi:10.1007/s12311-017-0870-9 (2017).

31. Edener, U. et al. Early onset and slow progression of SCA28, a rare dominant ataxia in a large four-generation family with a novel AFG3L2 mutation. Eur J Hum Genet 18, 965–968, doi:10.1038/ejhg.2010.40 (2010).

32. Cagnoli, C. et al. SCA28, a novel form of autosomal dominant cerebellar ataxia on chromosome 18p11.22-q11.2. Brain 129, 235–242, doi:10.1093/brain/awh651 (2006).

33. Charif, M. et al. A novel mutation of AFG3L2 might cause dominant optic atrophy in patients with mild intellectual disability. Frontiers in Genetics 6, doi:ARTN 311 10.3389/fgene.2015.00311 (2015).

34. Colavito, D. et al. Non-syndromic isolated dominant optic atrophy caused by the p.R468C mutation in the AFG3 like matrix AAA peptidase subunit 2 gene. Biomedical Reports 7, 451–454, doi:10.3892/br.2017.987 (2017).

35. Pierson, T. M. et al. Whole-Exome Sequencing Identifies Homozygous AFG3L2 Mutations in a Spastic Ataxia-Neuropathy Syndrome Linked to Mitochondrial m-AAA Proteases. Plos Genetics 7, doi:ARTN e1002325 10.1371/journal.pgen.1002325 (2011).

36. Eskandrani, A. et al. Recessive AFG3L2 Mutation Causes Progressive Microcephaly, Early Onset Seizures, Spasticity, and Basal Ganglia Involvement. Pediatric Neurology 71, 24–28, doi:10.1016/j.pediatrneuro1.2017.03.019 (2017).

37. Gates, S. N. et al. Ratchet-like polypeptide translocation mechanism of the AAA+ disaggregase Hsp104. Science 357, 273–279, doi:10.1126/science.aan1052 (2017).

38. Monroe, N., Han, H., Shen, P. S., Sundquist, W. I. & Hill, C. P. Structural basis of protein translocation by the Vps4-Vta1 AAA ATPase. Elife 6, doi:10.7554/eLife.24487 (2017).

39. Ripstein, Z. A., Huang, R., Augustyniak, R., Kay, L. E. & Rubinstein, J. L. Structure of a AAA plus unfoldase in the process of unfolding substrate. Elife 6 (2017).

40. de la Pena, A. H., Goodall, E. A., Gates, S. N., Lander, G. C. & Martin, A. Substrate-engaged 26S proteasome structures reveal mechanisms for ATP-hydrolysis-driven translocation. Science 362, 1018-+ (2018).

41. Yu, H. et al. ATP hydrolysis-coupled peptide translocation mechanism of Mycobacterium tuberculosis ClpB. Proc Natl Acad Sci USA 115, E9560–E9569, doi:10.1073/pnas.1810648115 (2018).

42. Ding, B. J., Martin, D. W., Rampello, A. J. & Glynn, S. E. Dissecting Substrate Specificities of the Mitochondrial AFG3L2 Protease. Biochemistry 57, 4225–4235, doi:10.1021/acs.biochem.8b00565 (2018).

43. Shi, H., Rampello, A. J. & Glynn, S. E. Engineered AAA plus proteases reveal principles of proteolysis at the mitochondrial inner membrane. Nature Communications 7, doi:ARTN 13301 10.1038/ncomms13301 (2016).

44. Rampello, A. J. & Glynn, S. E. Identification of a Degradation Signal Sequence within Substrates of the Mitochondrial i-AAA Protease. Journal of Molecular Biology 429, 873–885, doi:10.1016/j.jmb.2017.02.009 (2017).

45. Augustin, S. et al. An intersubunit signaling network coordinates ATP hydrolysis by m-AAA proteases. Mol Cell 35, 574–585, doi:10.1016/j.molcel.2009.07.018 (2009).

46. Tatsuta, T., Augustin, S., Nolden, M., Friedrichs, B. & Langer, T. m-AAA protease-driven membrane dislocation allows intramembrane cleavage by rhomboid in mitochondria. EMBO J 26, 325–335, doi:10.1038/sj.emboj.7601514 (2007).

47. Suno, R. et al. Structure of the whole cytosolic region of ATP-dependent protease FtsH. Molecular Cell 22, 575–585, doi:10.1016/j.molcel.2006.04.020 (2006).

48. Bieniossek, C., Niederhauser, B. & Baumann, U. M. The crystal structure of apo-FtsH reveals domain movements necessary for substrate unfolding and translocation. Proceedings of the National Academy of Sciences of the United States of America 106, 21579–21584, doi:10.1073/pnas.0910708106 (2009).

49. Vostrukhina, M. et al. The structure of Aquifex aeolicus FtsH in the ADP-bound state reveals a C-2-symmetric hexamer. Acta Crystallographica Section D-Structural Biology 71, 1307–1318, doi:10.1107/S1399004715005945 (2015).

50. Wendler, P., Ciniawsky, S., Kock, M. & Kube, S. Structure and function of the AAA+ nucleotide binding pocket. Biochim Biophys Acta 1823, 2–14, doi:10.1016/j.bbamcr.2011.06.014 (2012).

51. Karata, K., Inagawa, T., Wilkinson, A. J., Tatsuta, T. & Ogura, T. Dissecting the role of a conserved motif (the second region of homology) in the AAA family of ATPases. Site-directed mutagenesis of the ATP-dependent protease FtsH. J Biol Chem 274, 26225–26232 (1999).

52. Akiyama, Y. Self-processing of FtsH and its implication for the cleavage specificity of this protease. Biochemistry 38, 11693–11699, doi:DOI 10.1021/bi991177c (1999).

53. Graef, M., Seewald, G. & Langer, T. Substrate recognition by AAA+ ATPases: distinct substrate binding modes in ATP-dependent protease Yme1 of the mitochondrial intermembrane space. Mol Cell Biol 27, 2476–2485, doi:10.1128/MCB.01721-06 (2007).

54. Herzik, M. A., Wu, M. Y. & Lander, G. C. Achieving better-than-3-angstrom resolution by single-particle cryo-EM at 200 keV. Nat Methods 14, 1075–1078 (2017).

55. Suloway, C. et al. Automated molecular microscopy: the new Leginon system. J Struct Biol 151, 41–60, doi:10.1016/j.jsb.2005.03.010 (2005).

56. Lander, G. C. et al. Appion: an integrated, database-driven pipeline to facilitate EM image processing. J Struct Biol 166, 95–102 (2009).

57. Zheng, S. Q. et al. MotionCor2: anisotropic correction of beam-induced motion for improved cryo-electron microscopy. Nat Methods 14, 331–332, doi:10.1038/nmeth.4193 (2017).

58. Rohou, A. & Grigorieff, N. CTFFIND4: Fast and accurate defocus estimation from electron micrographs. J Struct Biol 192, 216–221, doi:10.1016/j.jsb.2015.08.008 (2015).

59. Roseman, A. M. FindEM–a fast, efficient program for automatic selection of particles from electron micrographs. J Struct Biol 145, 91–99 (2004).

60. Scheres, S. H. RELION: implementation of a Bayesian approach to cryo-EM structure determination. J Struct Biol 180, 519–530, doi:10.1016/j.jsb.2012.09.006 (2012).

61. Zivanov, J. et al. New tools for automated high-resolution cryo-EM structure determination in RELION-3. Elife 7, doi:10.7554/eLife.42166 (2018).

62. Heymann, J. B. Guidelines for using Bsoft for high resolution reconstruction and validation of biomolecular structures from electron micrographs. Protein Sci 27, 159–171, doi:10.1002/pro.3293 (2018).

63. Goddard, T. D., Huang, C. C. & Ferrin, T. E. Visualizing density maps with UCSF Chimera. J Struct Biol 157, 281–287, doi:10.1016/j.jsb.2006.06.010 (2007).

64. Emsley, P. & Cowtan, K. Coot: model-building tools for molecular graphics. Acta Crystallogr D Biol Crystallogr 60, 2126–2132, doi:10.1107/S0907444904019158 (2004).

65. Afonine, P. V. et al. Towards automated crystallographic structure refinement with phenix.refine. Acta Crystallogr D Biol Crystallogr 68, 352–367, doi:10.1107/S0907444912001308 (2012).

66. Herzik, M. A., Jr., Fraser, J. S. & Lander, G. C. A Multi-model Approach to Assessing Local and Global Cryo-EM Map Quality. Structure 27, 344–358, doi:10.1016/j.str.2018.10.003 (2019).

67. Haynes, C. M., Yang, Y., Blais, S. P., Neubert, T. A. & Ron, D. The matrix peptide exporter HAF-1 signals a mitochondrial UPR by activating the transcription factor ZC376.7 in C. elegans. Mol Cell 37, 529–540, doi:10.1016/j.molcel.2010.01.015 (2010).

68. Geissler, A. et al. The mitochondrial presequence translocase: an essential role of Tim50 in directing preproteins to the import channel. Cell 111, 507–518 (2002).

69. Barad, B. A. et al. EMRinger: side chain-directed model and map validation for 3D cryo-electron microscopy. Nat Methods 12, 943–946, doi:10.1038/nmeth.3541 (2015).

